# Single-probe, high-yield spatial mapping of diverse short RNA motifs

**DOI:** 10.64898/2026.11.02.763225

**Authors:** Guohao Liang, Kenneth Joseph, Kayleigh Sim, Vadym Kuchumov, An Doan, Zhiqun Tan, Xiangmin Xu, Fangyuan Ding

**Author notes:** These authors contributed equally (Guohao Liang, Kenneth Joseph).

## Abstract

Spatial transcriptomics has revolutionized our understanding of RNA biology. Hybridization-based methods map mRNA transcripts by tiling multiple labeled probes along each molecule to generate a visible fluorescent signal, restricting applicability to long transcripts. Short RNA motifs, consequently, can only accommodate a single probe and remain a spatially invisible dimension of RNA biology. Here we present SMOOTHY-FISH, a single probe strategy that directly visualizes short RNA motifs at the single-molecule resolution. By optimizing probe brightness, binding specificity, and target accessibility, our single ∼20-nucleotide probe reached the practical ceiling of hybridization-based detection. We quantitatively mapped previously inaccessible short motifs — splice-isoform junctions, individual microRNAs, and structural conformations — in single cells and mouse brain tissues, each validated against orthogonal methods. This unveiled biological phenomena that were previously unknown: ratios of alternatively spliced isoforms are tightly regulated across cell populations, subcellular miRNA fractions are precisely maintained, and a designed RNA structure was found to have a prominent alternative conformation. These findings demonstrate that short RNA motifs possess key spatial information that has yet to be discovered. With SMOOTHY-FISH’s optimized single probe technique, short RNA motifs are converted into a new routinely measurable axis of spatial transcriptomics.

## Main

Expanding the scope of spatial transcriptomics has progressively revealed new information about RNA biology^1^. Single-molecule fluorescence in-situ hybridization (smFISH) first enabled the direct visualization and quantification of individual mRNA molecules with subcellular resolution, revealing RNA localization and transcriptional heterogeneity between cell types^2,3^. Barcoding techniques have extended these capabilities to hundreds or thousands of RNA species simultaneously, enabling system-wide analysis of gene expression, cell identity, and tissue organization^4–6^. In-situ sequencing methods have further enabled untargeted, transcriptome-side profiling of RNA species directly within intact tissues^7–10^. Together, these improvements have established spatial RNA imaging as a powerful approach for quantifying and localizing messenger RNA within their native cellular context, revealing insights into cell fates, cell-to-cell interactions, and gene activity across complex tissues^1^.

Despite these advancements, these methods rely on the cooperative hybridization of multiple probes along a single molecule to generate sufficient signal for detection^2–8^. This requirement restricts their application to RNAs long enough to contain hundreds of nucleotides of unique, tileable sequences, applying mostly to messenger RNA (mRNA) (Fig. 1a). Reducing the number of probes can be achieved by introducing signal amplification at the cost of single-molecule quantification and yield^11–13^. Many biologically important RNA features, however, are defined by much shorter motifs than current methodologies can accommodate. These include but are not limited to short sequence-specific motifs embedded within long transcripts, such as isoform-defining junctions^14,15^; short RNA molecules containing roughly 20-nucleotides (nt) long, such as mature miRNAs^16–19^; or RNA structures that contain a short-junction motif generated by non-linear RNA folding (Fig. 1a)^20–22^. These features have remained indiscernible to spatial imaging not because they are unimportant, but simply because they are too short.

**Figure 1.** A single-probe strategy—SMOOTHY-FISH—expanded the scope of single-molecule spatial RNA mapping. **a**, The scope of single-molecule spatial RNA mapping is constrained by both the RNA molecular length and the target-specific motif length. Current tools relying on multiple probes cover only long motifs residing in long RNAs (mRNAs) while our single-probe strategy — SMOOTHY-FISH — extends the coverage to short motifs in both short RNAs (small RNAs) and long RNAs with short motifs (RNA structures, RNA isoforms). **b**, A reliable single-probe strategy needs to satisfy three requirements: probe design to enhance signal brightness, binding specificity to reduce non-specific or off-targeting signals, and target accessibility optimization to maximize detection yield. **c**, SMOOTHY-FISH’s detection yield approached the practical ceiling set by the smFISH benchmark. ① SMOOTHY-FISH’s detection yield was measured through colocalization between SMOOTHY-FISH (green) and smFISH (red) on SDHA mRNAs. The benchmarks were measured through colocalization between two sets of smFISH on SDHA mRNAs: ② upstream (green) and downstream (red) probe set; ③ odd (green) and even (red) probe set. Sequences of SMOOTHY-FISH and smFISH probes are listed in Supplementary Table 1, 2, & 5. In the image of SDHA mapping in an HEK293 cell in ①, fluorescent signals corresponding to smFISH (red) and SMOOTHY-FISH (green) are shown in zoom insets, with colocalized dots circled in orange and non-colocalized dots circled in blue. Detection yields in ①, ②, ③ are shown as mean ± SD (50 cells). The benchmark is shown as mean (dash line) ± SD (gray zone). See more on detection yield in Supplementary Note 1.

Mapping short RNA motifs requires a detection strategy that differs fundamentally from imaging a long transcript. Limited unique sequences can only accommodate a singular probe that must have a detection yield comparable to the combination of tens of probes in conventional smFISH. This imposes three key requirements (Fig. 1b): (1) the probe must generate a signal bright enough on its own to be quantifiable as an individual molecule while remaining distinguishable from cellular background; (2) it must be highly specific as a single non-specific probe will be indistinguishable from a true target-motif signal; (3) the target motif must be accessible to the single probe since there are no additional binding sites to compensate for a failed hybridization. Meeting all three requirements simultaneously with a single probe remains a challenge that has kept short motifs inaccessible to spatial transcriptomics.

Here we present **SMOOTHY-FISH** (**S**hort-motif **M**inimum **O**f **O**ff-**T**arget **H**igh-**Y**ield FISH), a modular single-probe imaging strategy that satisfies all three requirements and enables robust detection of short RNA motifs in cells and intact tissues. By optimizing probe design, binding specificity, and target accessibility, a single ∼20-nt probe detects its target as reliably as a full set of tiled smFISH probes against the same transcript, reaching the same practical yield with a fraction of the sequence (Fig. 1c). Using a modular single-probe, we quantitatively mapped three classes of short-motifs that were previously unresolvable — splice-isoform junctions, individual microRNAs, and RNA structural conformations — each validated against orthogonal measurements.

### Optimization of SMOOTHY-FISH probes and single-probe workflow

Guided by the signal, specificity, and accessibility requirements, we first established a standard for our single-probe to be measured against. Because no hybridization-based method is perfect, the highest achievable yield cannot reach 100%^23^. We determined a practical benchmark by utilizing two independent smFISH probe sets against the same target (Fig. 1c; Supplementary Note 1). Each set contained 21 probes and were simultaneously hybridized to the endogenous SDHA mRNA in HEK293 cells (Supplementary Table 1, 2). We observed detection yields through colocalization of 69.94±5.96% and 73.75±5.82% for separated and alternating probe sets respectively (Fig. 1c.; Supplementary Note 1). This colocalization readout provides the highest practical detection yield for hybridization-based single-molecule imaging, and thus the benchmark a single probe design should approach. Through probe design, binding specificity, and accessibility optimizations described below, SMOOTHY-FISH reaches this practical benchmark using a single ∼20-nt probe (Fig. 1c).

### Probe Design

We designed SMOOTHY-FISH probes with a modular architecture that separates target recognition from signal generation. Each SMOOTHY-FISH probe consists of three modules: (i) A short, 20-nt single-stranded DNA region to bind the target motif; (ii) a fluorophore-bearing double-stranded DNA region; (iii) and a flexible linker connecting these two functional regions to reduce steric hindrance (Fig. 2a). This design allows SMOOTHY-FISH to incorporate multiple fluorescent dyes for signal generation while keeping the binding region sufficiently short for its target’s motif.

**Figure 2.** Optimization of SMOOTHY-FISH probe design and binding specificity. **a**, The SMOOTHY-FISH probe design consists of three modules: a ∼ 20-nt single-stranded target-binding region, a double-stranded signal generation region bearing multiple fluorophores, and a flexible linker to avoid steric hindrance in hybridization. **b**, SMOOTHY-FISH probes were optimized by altering the combination of fluorophore number (n) and the gap size (nt) between neighboring fluorophores. The optimal combination is represented in red in the graph. Images of suboptimal probe designs can be found in Supplementary Fig. 1. Brightness quantification can be found in Supplementary Table 3, Supplementary Note 1 and 2. **c**, Detection yield comparison among 5-nt gap probes showed 5 fluorophores with 5-nt gap achieved the highest detection yield. Detection yields are represented by mean ± SD (n = 50 cells). Notably, detection yield dropped drastically with 8 or more fluorophores on the probes. **d**, The optimal design (n = 5; gap = 5nts) was compatible with various fluorophores. We mapped SDHA mRNA by a single SMOOTHY-FISH probe with Alexa Fluor 488 (green), Alexa Fluor 594 (yellow), and ATTO647N (red). **e**, SMOOTHY-FISH’s small RNAs retention in hydrogel. We verified that small RNA oligonucleotides could be poly(A)-tailed by E.coli poly(A) polymerase. PAGE left lane: negative control without E.coli poly(A) polymerase; right lane: poly(A)-tailing on a 24-nt RNA oligonucleotide. After in situ poly(A) tailing, 5’-poly(dT) oligonucleotides were applied to hybridize to poly(A) tails, followed by polyacrylamide hydrogel embedding and clearing to remove proteins and lipids. **f**, SMOOTHY-FISH signals were linearly correlated to the transfection concentration when targeting the transfected 24-nt orthogonal RNA oligonucleotides in HEK293 cells. SMOOTHY-FISH signals were represented by mean ± SD (n = 50 cells). Sequences of the orthogonal RNA oligonucleotide and the corresponding probe used in **e** and **f** are stated in Supplementary Table 4.

### Signal

To generate a bright single-molecule signal, we optimized the trade-off between brightness and probe size by modifying two parameters that govern the fluorophore-bearing region (Fig. 2b). The first parameter we systematically compared was the spacing between fluorophores. Previous multi-labeled probe designs often space out their fluorophores widely due to known self-quenching effects and other fluorophore-fluorophore interactions at close proximity (Fig. 2b)^2,24^. Increasing the spacing between the fluorophores reduces the quenching effect, but in-turn causes the probe size to increase dramatically (Fig. 2b). Finding a balance between fluorophore quenching and probe diffusion is critical for SMOOTHY-FISH probes to reach its target and produce a visible signal (Fig. 2b). Remarkably, we found that a relatively short distance of 5-nt spacing was sufficiently wide for SMOOTHY-FISH probes to generate bright, single-molecule signals (Fig. 2c; Supplementary Note 2)^24^. Further reducing the spacing to 4 or 3-nt decreased the probe brightness, consistent with quenching-associated signal loss (Supplementary Fig. 1a; Supplementary Note 2).

We then refined the second parameter: the number of fluorophores carried by each probe. While the fluorophore-bearing was made double-stranded to minimize unintended hybridization to cellular RNAs, each additional fluorophore further increases probe size and in turn may reduce probe diffusion and target hybridization (Fig. 2b). To assess this balance, we quantified the colocalization rate of probes with different numbers of fluorophores spaced 5-nt apart. As expected, utilizing 1 or 2 fluorophores resulted in signals indistinguishable from background noise (Supplementary Fig. 1b). Probes carrying 8 or more fluorophores, corresponding to duplexes longer than 63-nt, showed a substantial decrease in yield, suggesting a reduction in probe diffusion or hybridization efficiency (Fig. 2c; Supplementary Fig. 1c; Supplementary Note 2). While 3 or 4 fluorophores produced visible signals, 5 fluorophores corresponding with a 45-nt duplex showed the highest yield, maximizing the balance of probe size and signal generation (Fig. 2c). Across all three fluorophores tested: Alexa-488, Alexa-594, and ATTO-647N, the 5 fluorophore 5-nt spacing probes produced robust, quantifiable single-molecule dots, and thus were used for all subsequent experiments (Fig. 2d; Supplementary Table 3, Supplementary Note 2).

### Specificity

Since a singular probe carries the entire detection burden, any non-specifically retained probes will produce a signal indistinguishable from a true one. We therefore adopted a hydrogel-based RNA anchoring and cell clearing workflow to increase our specificity^25^. This technique is used to: (i) preserve the spatial location of our target RNAs, (ii) remove unwanted cellular components including proteins and lipids to reduce non-specific probe retention and molecular crowding, and (iii) increase SMOOTHY FISH’s ability to diffuse across a cleared sample for hybridization. Previous methods accomplish this by hybridizing a modified oligonucleotide to the poly(A) tail of mRNAs to cross link them to the hydrogel^5^.

While hydrogel-based anchoring is well established for mRNAs, non-coding RNAs lack a poly(A) to be targeted in the same manner. To expand upon this method, we utilized a template-independent *E. coli* polyA polymerase to add an artificial polyA tail to non-coding RNAs to also retain them within the hydrogel (Fig. 2e)^26,27^. In-vitro tests showed that a 24-nt RNA substrate can be extended by more than 150 adenosines to the 3’ end after 16 hours (Fig. 2e). This extended tail allows for multiple previously used poly(dT) anchors to hybridize on the artificial tail, efficiently retaining short non-coding RNAs in the hydrogel system for SMOOTHY-FISH mapping.

Given that the short length of small RNAs precludes colocalization-based validation, we quantitatively assessed the performance of SMOOTHY-FISH’s capture of non-coding short RNAs. We transfected a miRNA mimic into HEK293 cells, and compared the concentration of the miRNA with the total number of SMOOTHY-FISH dots (Supplementary Table 4). Across input concentrations from 1 to 5 nM, SMOOTHY-FISH mapping showed a linear increase in captured RNA from 12.26 to 98.46 dots per cell, confirming the proper capture of small RNAs (Fig. 2f). Without any transfection of the miRNA mimic, SMOOTHY-FISH probes showed minimal off-target hybridization with only 0.84 dots per cell (Fig. 2f, Supplementary Fig. 2a, b). In addition, the control samples processed without the poly(A) tailing showed fewer than 1.24 dots per cell across all concentrations (Fig. 2f). These results demonstrate that an in-situ poly(A) tailing is necessary for short RNA retention and enables SMOOTHY-FISH to quantitatively report short motif abundance.

### Accessibility

With signal and specificity optimized, the last requirement to be met is the accessibility of a single probe to the target. To improve the detection efficiency of SMOOTHY-FISH towards our practical benchmark, we sought to increase the local accessibility of the target site by utilizing a pair of 40-nt flanking oligonucleotides (Fig. 3a)^28^. The native structure of the RNA can reduce the ability of SMOOTHY-FISH probes to bind, causing variability in the detection efficiency depending on the chosen target sequence across the transcript^29,30^. With small motifs limiting the possible binding regions, we hypothesized that flanking oligonucleotides would help relax local secondary structures and allow SMOOTHY-FISH probes to access their target^28^.

**Figure 3.** Optimization of SMOOTHY-FISH target accessibility by flanking oligonucleotides. **a**, We hybridized a pair of flanking oligonucleotides next to the target site to optimize the target accessibility for SMOOTHY-FISH probe hybridization. **b**, SMOOTHY-FISH mapping of site P07, P17, and P26 on SDHA mRNA showed that flankers greatly increased the detection yields. Comparison among the distance between flankers and target site showed that detection yield increase was positively correlated with the proximity in less-accessible sites (P07, P17). Detection yields were represented by mean ± SD (n = 50 cells). Sequences of smFISH probes, SMOOTHY-FISH probes, and flankers are listed in Supplementary Table 1, 2, & 5. **c**, Flankers immediately adjacent to site P17 maximized the colocalization between SMOOTHY-FISH (green) and smFISH (red) in SDHA mRNA mapping. Colocalized dots are circled in yellow; non-colocalized dots are circled in blue. **d**. SMOOTHY-FISH (green) mapping of mmuSDHA site P12 showed high colocalization to smFISH (red) in mouse brain cortex. Sequences of SMOOTHY-FISH probes, smFISH probes, and flankers are listed in Supplementary Table 6.

To check for improvements in local accessibility, we chose three different binding regions across the endogenous human SDHA: P07, P17, and P26 (Fig. 3b; Supplementary Table 5). Without any flanking oligonucleotides, the detection efficiencies for each were 35.87±8.98%, 50.65±8.49%, and 69.57±5.44% respectively, showing high variability of accessibility across the SDHA mRNA (Fig. 3b; Supplementary Note 1)^31^. While most of the SMOOTHY-FISH dots colocalize with smFISH, a significant amount of SDHA mRNA remains without a SMOOTHY-FISH signal for regions with lower accessibility (Fig. 3c). We then designed pairs of 40-nt flankers to hybridize 0 to 4-nt away from the target site on the mRNA to test how close the flankers need to be to improve accessibility (Supplementary Table 5). For P17, adding flanking oligonucleotides 4-nt away from the target site increased detection efficiency to 60.25±5.40%, reaching as high as 67.68±6.06% when placed directly next to the target (Fig. 3b). With the improvements to target accessibility, SMOOTHY-FISH performed as well as smFISH in detecting SDHA mRNA at the P17 site (Fig. 3c). Similarly to P17, P07 also saw significant improvements with the addition of adjacent flankers (Figure 3B). P26, on the other hand, saw limited improvements as its yield was already comparable to the practical benchmark, indicating that the site was already accessible (Fig. 1c, 3b).

Finally, we evaluated SMOOTHY-FISH’s within the mouse brain to ensure compatibility with intact tissues. We targeted the endogenous mouse SDHA at the mmuP12 position which contained the same sequence as the human P26 (Supplementary Table 6). When tested in the cortex after introduction of the hydrogel and flanker system, mmuP12 detection achieved an average yield of 58.55%, despite elevated tissue background (Fig. 3d; Supplementary Note 1). These results indicate that SMOOTHY-FISH maintains efficient detection in tissue samples and can support RNA mapping in complex environments.

### Mapping important but previously inaccessible RNA targets using SMOOTHY-FISH

Having cleared the three requirements and establishing SMOOTHY-FISH as a reliable single-probe method, we can now investigate the regions of RNA biology that were previously inaccessible by long-transcript methods (Fig. 1a). Targeting short-motifs is not one problem but several, such as isoform junctions embedded in long transcripts, small RNAs shorter than conventional multi-probe targets, and structural junctions formed by RNA folding. We applied the same modular SMOOTHY-FISH design to all three cases to overcome their detection challenges: discriminating near-identical sequences, capturing molecules too short for any redundancy, and reading out sequences generated by structure rather than by the genome.

### Splicing isoforms

More than 95% of human genes undergo alternative splicing, where exons are selectively combined to generate distinct RNA isoforms from a single gene^32^. The inclusion or exclusion of different exons can change protein function, localization, stability, or translation efficiency depending on its location within the transcript^14^. A well-characterized example is pyruvate kinase M (PKM), which produces two isoforms PKM1 and PKM2 through the inclusion of exon 9 or exon 10, respectively (Supplementary Note 3)^33,34^. PKM1 is constitutively expressed in tissues with high metabolic demands, such as the brain or skeletal muscle, while PKM2 is dynamically regulated and promotes biosynthetic pathways in proliferating cells and tumors^34,35^. Despite their distinct functional properties, isoforms typically exhibit high sequence similarity, with the specific exon-exon junction of PKM1 and PKM2 only differing by 2-nt at the third and sixth positions (Fig. 4a). This limited sequence divergence poses a significant challenge for accurate isoform-specific detection using most hybridization-based methods.

**Figure 4.** SMOOTHY-FISH revealed a tight expression ratio between PKM1 and PKM2 isoforms despite a wide range of expression levels in cell lines and in mouse brain tissue. **a**, Alternative splicing of PKM generates two isoforms PKM1 (blue, containing exon 9) and PKM2 (orange, containing exon 10) that differ by only two nucleotides. **b**, SMOOTHY-FISH probe targeted the PKM1 (green) and PKM2 (red) spliced exon junction. **c**, SMOOTHY-FISH distinguished PKM1 (green) and PKM2 (red) in HEK293 cells. Notably, PKM1 and PKM2 dot signals did not overlap, showing the high specificity of SMOOTHY-FISH. Zoomed image insets showed PKM2 was predominantly expressed in cells with different total PKM expression levels. **d**, The expression levels of PKM1 and PKM2 exhibited high variation within the cell population. PKM1 and PKM2 dot counts per cell were represented by mean ± SD in the bar graph (n = 50 cells). **e**, The ratio of Pkm1 (green) and Pkm2 (red) was flipped in the mouse hippocampus compared to cultured cell lines. Zoomed image inset showed Pkm1 was predominantly expressed across different regions in the mouse hippocampus. Sequences of SMOOTHY-FISH probes and flankers are listed in Supplementary Table 7. **f**. PKM1 and PKM2 expression ratios were very tight despite high variation in the total PKM expression across individual cells. The PKM2 fractions were estimated as the slopes of the linear fitting. The tight linear correlations are represented by the R^2^ from the linear fitting. See the PKM2 fraction calculation in Supplementary Note 3.

SMOOTHY-FISH provides a practical approach for distinguishing transcript isoforms by targeting the isoform-specific splice junctions with its improved single probe design. We created two distinctly labeled SMOOTHY-FISH probes targeting the 8/9 exon junction for PKM1 and the 8/10 exon junction for PKM2 (Fig. 4b; Supplementary Table 7). The two probe signals had no observable colocalization, consistent with the mutual exclusivity of the alternatively spliced exons (Fig. 4c, Supplementary Fig. 3). To ensure the fluorescent dyes themselves do not contain any bias, we repeated the experiment with the dyes reversed and saw no observable differences (Supplementary Fig. 4; Supplementary Table 7).

In both HEK293 cells and SH-SY5Y cells, PKM2 had a significantly higher transcript count than PKM1 (Fig. 4c, d). For HEK293 cells, average transcript counts for PKM1 and PKM2 were 25.60 ± 13.75 and 156.56 ± 40.10 respectively, yielding an average ratio of PKM1 to PKM2 of 1:6.09 (Fig. 4d). Similarly, SH-SY5Y cells had counts of 12.72 ± 7.35 and 122.60 ± 61.60, yielding a PKM1 to PKM2 ratio of 1:9.64 (Fig. 4d). We validated SMOOTHY-FISH’s quantitative measurements of the isoform ratios by using RT-PCR and restriction digestion, finding the fraction of PKM2 isoforms to between 1:6.66 for HEK293 cells and 1:9.85 for SH-SY5Y cells respectively, confirming our SMOOTHY-FISH results (Supplementary. Fig. 5a, b; Supplementary Note 3)^36^.

While immortalized cells are focused on rapid cell growth and division, neurons within the brain are consuming high amounts of energy, requiring more of the PKM1 isoform. Consistent with previous studies of neurons and the brain, expression levels of PKM1 were higher than PKM2 within the hippocampus (Fig. 4e; Supplementary Table 7)^34^. SMOOTHY-FISH quantification of the ratio of PKM1 to PKM2 in the hippocampus was 2.84:1, with confirmation by RT-PCR and restriction digestion yielding a similar ratio of 3.41:1 (Supplementary Fig. 5c; Supplementary Note 3; Supplementary Table 7). These results highlight the ability for SMOOTHY-FISH to accurately distinguish between PKM1 and PKM2 based on their splicing junctions.

Because SMOOTHY-FISH counts isoforms one molecule at a time, it also reports how the isoform ratio is distributed across single cells. Despite having high cell-to-cell heterogeneity of isoform transcript abundance, the individual ratio of cells remained remarkably consistent (Fig. 4b, d). The distribution of the ratio across cells was unexpectedly tight, with the ratio of PKM1 to PKM2 of 1:5.85 in HEK293 cells and 1:9.75 in SH-SY5Y cells, with R^2^ values of 0.946 and 0.995 respectively (Fig. 4f; Supplementary Fig. 6; Supplementary Note 3). Despite the large variation in expression, the defined ratio suggests that cells tightly regulate the PKM gene to properly balance the isoforms at a single-cell level, illustrating an observation that can only be found once isoforms are individually counted in-situ (Supplementary Note 3).

### Small RNAs

MiRNAs are an indispensable class of small regulatory RNAs that modulate gene expression through sequence-specific repression of target mRNAs^17–19^. The expression patterns of miRNAs depend on the cell type and their specific purpose, with most localizing within the cytoplasm^37^. Despite extensive studies revealing miRNA functions, their 20 to 25-nt size presents a significant challenge for spatial visualization. Although sequencing and stem-loop real-time PCR have been widely used to profile the relative expression level across miRNAs, methods to resolve individual miRNA molecules in intact cells and tissues and to quantify the absolute miRNA expression remain to be seen^38,39^.

Similarly to the validation of short non-coding RNAs in figure 2f, colocalization-based validation is not feasible for miRNAs since the binding sequence can only accommodate a single probe. We therefore validated SMOOTHY-FISH’s ability to capture and spatially resolve miRNAs by choosing three endogenous miRNAs in HeLa cells with increasing expression levels: miRNA-10a-5p, miRNA-16-5p, and miRNA-23a-3p (Supplementary Note 4)^40^. SMOOTHY-FISH mapping of all three miRNAs revealed 58.20 ± 19.84, 145.10 ± 58.18, and 270.16 ± 75.69 dots per cell respectively, matching previously reported relative expression levels (Fig. 5a; Supplementary Fig. 7a - d, Supplementary Table 8)^40^. The polyA-tailing method was again necessary for reliable retention, identification, and mapping of all three miRNAs (Fig. 5a).

**Figure 5.** SMOOTHY-FISH demonstrated that miRNAs predominantly localized to cytoplasm at tight fractions albeit the expression variation across individual cells. **a**, SMOOTHY-FISH was able to map miRNAs with a wide range of abundance in cells. miR-10a-5p, miR-16-5p, miR-23a-3p were reported to have low, intermediate, and high expression in HeLa cells respectively. SMOOTHY-FISH mapped these three miRNAs with and without the poly(A) tailing process. miRNA dot counts per cell are represented by mean ± SD in the bar graph (n = 50 cells). SMOOTHY-FISH mapped hippocampus-enriched miR-218-5p in **b**, hippocampus, **c**, midbrain, and **d**, hypothalamus of the mouse brain tissue with and without poly(A) tailing. **e**, miR-218-5p predominantly localized to cytoplasm in 3T3 cells (n = 50 cells). poly(A) tailing was required to retain and map miRNAs in cell lines and in mouse brain tissues. **f**, All four mapped miRNAs localized to cytoplasm at tight fractions despite the expression variation across individual cells (n = 50 cells). Linear fitting was performed between total miRNAs and cytoplasmic miRNAs for miR-10a-5p, miR-16-5p, miR-23a-3p, miR-218-5p in cell lines. The tight linear correlations are represented by the R^2^ from the linear fitting. See the PKM2 fraction calculation in Supplementary Note 4. Sequences of SMOOTHY-FISH probes for miRNA mapping are listed in Supplementary Table 8.

With tissue samples containing complex environments and cell type heterogeneity, we validated SMOOTHY-FISH’s ability to capture and spatially resolve miRNAs within mouse brain tissue samples by choosing a region specific miRNA, miRNA-218-5p (SI Note 4)^41^. MiRNA-218-5p has been reported to be highly enriched in the hippocampus, where it plays important roles in neuronal circuit assembly and cognitive function (Supplementary Fig. 8)^42^. SMOOTHY-FISH detection of miRNA-218-5p in the mouse brain revealed high expression levels within the hippocampus, especially in highly dense cellular regions such as the cornu ammonia (Fig. 5B, Supplementary Table 8). As previously exhibited in cells, poly(A)-tailing was still necessary for miRNA detection in tissue samples despite the dense tissue architecture (Fig. 5b). In other brain regions, such as the midbrain and hypothalamus, miRNA-218-5p was expressed substantially lower than observed in the hippocampus, demonstrating the region-specificity of this miRNA as seen in prior observations (Fig. 5c, d).

SMOOTHY-FISH’s mapping of miRNAs extends beyond simply mapping their expression levels; we can also visualize miRNA patterns at a subcellular level. For miRNA-218-5p within the hippocampus, approximately 82.6% of the miRNA was found to localize within the cytoplasm (Supplementary Note 4). This cytoplasmic ratio matches the miRNA-218-5p patterns found in 3T3 cells, where 87.3% localized within the cytoplasm (Fig. 5e, f). The three miRNAs tested in HeLa cells also favored cytoplasmic localization with 74.5% for miRNA-10a-5p, 77.5% for miRNA-16-5p, and 68.0% for miRNA-23a-3p (Fig. 5f). Each miRNA displayed a higher cytoplasmic fraction, consistent with miRNAs functioning within the cytoplasm by post-transcriptional gene silencing^37^.

Furthermore, the cytoplasmic fraction appeared remarkably consistent across single cell populations, despite a larger variance in the total expression of each miRNA. We observed a strong linear relationship between individual miRNA abundance and their amounts localized within the cytoplasm. The R^2^ values for cytoplasmic abundance were 0.903 for miRNA-10a-5p, 0.958 for miRNA-16-5p, 0.811 for miRNA-23a-3p, and 0.957 for miRNA-218-5p (Fig. 5f). This indicates that cells tightly control the mechanisms of cytoplasmic and nuclear localization to ensure that miRNA targets are properly regulated in both compartments (Supplementary Note 4).

With the subcellular fractions of miRNAs strongly controlled, we reasoned that variability in miRNA abundance could be due to cellular size. We therefore explored the nuclear and cytoplasmic concentrations, indicated by the relationship between subcellular abundance and area. The nuclear concentration of each miRNA showed a negligible relationship (Supplementary Fig. 9a, Supplementary Note 4). Conversely, the cytoplasmic concentration showed a linear correlation with R^2^ values ranging from 0.541 to 0.727 across all four miRNAs (Supplementary Fig. 9b; Supplementary Note 4). The regulation of cytoplasmic concentration, but not nuclear concentration, is indicative of miRNA suppression of mRNAs, which are also regulated to maintain their cellular concentration (Supplementary Note 4)^43^.

### RNA structural motifs

RNAs encode function not only in their nucleotide sequences but also in the structures they form. By organizing functional regions and regulating their accessibility, RNA structures can influence gene expression at multiple stages, including transcription, RNA processing, translation, and degradation^20–22^. These structural features can further modulate the accessibility of regulatory elements and binding sites, influencing RNA stability, subcellular localization, and interactions with cellular factors^20–22^. Despite the fundamental importance of RNA structures within these processes, it has remained challenging to directly map RNA structures within their native spatial contexts^44^. A folded structure creates a defining feature that SMOOTHY-FISH is suited to detect: a junction that is formed by sequences that are distant in the primary transcript but brought together only when a structure is formed.

To demonstrate SMOOTHY-FISH’s ability to map RNA structures, we selected a well-characterized synthetic RNA structural element MS2-3-PP7-NiNTA as a model target (Supplementary Note 5)^45^. This 69-nt RNA comprises four structured modules: a 4-nt basal stem, MS2 and PP7 hairpins, and a NiNTA aptamer hairpin (Fig. 6a)^45^. The MS2 and PP7 aptamers are strategically positioned to enable bioluminescent tracking of RNA dynamics in live cells through the recruitment of complementary NanoLuc fragments, with the NiNTA aptamer added to lock the structure and allow for the RNA to be captured for downstream analysis (Fig. 6a)^45^. The fully folded MS2-3-PP7-NiNTA contains a structurally constrained junction across the transcript (Fig. 6a)^30^.

**Figure 6.** SMOOTHY-FISH revealed an unexpected conformation of the synthetic RNA structure element MS2-3-PP7-NiNTA in its native cellular environment. **a**, The synthetic RNA structural element MS2-3-PP7-NiNTA, where the basal stem (orange), MS2 hairpin (pink), PP7 hairpin (yellow), and NiNTA hairpin (blue) all folded, is the most stable conformation predicted in silico (Conformation *I*). **b**, Two-colored mapping was performed by targeting a common region 5’ upstream of the structural element and the Conformation *I* specific junction. SMOOTHY-FISH showed many, but not all, colocalization between the common region signals (green) and the Conformation *I* specific junction signals. Colocalized dots are circled in the zoomed image inset. The non-colocalized common region dots (green) suggested the possibility of other conformations. See more about how the junction probe sequences were designed in Supplementary Note 5. **c**, Table summary of the seven possible conformations mapped by SMOOTHY-FISH. These conformations were defined by different combinations of folded and unfolded states of the basal stem (orange), MS2 hairpin (pink), PP7 (hairpin), and NiNTA hairpin (blue). See the details of these seven conformations in Supplementary Fig. 11 and Supplementary Note 5. **d**, SMOOTHY-FISH revealed the co-existence between the Conformation *I* and Conformation *V*. Left: design of the SMOOTHY-FISH probe targeting the Conformation *V* junction (cyan). Right: three-colored SMOOTHY-FISH mapping of the common region (green), Conformation *I* junction (red), and Conformation *V* junction (cyan). Colocalized common-region and Conformation *I* dots are marked with circles and colocalized common-region and Conformation *V* dots are marked with squares in the zoomed image insets. See the three-colored SMOOTHY-FISH mapping of the common region, Conformation *I* junction, other conformation junctions in Supplementary Fig. 12. Sequences of probes for conformation mapping are listed in Supplementary Table 9. **e**, Colocalization fraction quantification of the three-colored mapping results showed Conformation *I* and Conformation *V* were the two co-existed and dominant conformations of the MS2-3-PP7-NiNTA structure element in its native cellular environment. Colocalization fractions with the common region were obtained by three-color mapping of the common region, the Conformation *I* junction, and each junction from the Conformation *II* – *VII* junctions in each experiment. Colocalization fractions are shown as medians with the 12.5^th^–87.5^th^ percentile ranges (n = 50 cells) in the bar graph. See the calculation in Supplementary Note 5.

To visualize the existence of the MS2-3-PP7-NiNTA structural element, we designed one SMOOTHY-FISH probe across this junction, with 10-nt on each side, and a second probe targeting a 20-nt region outside the structure as a reference for total transcript abundance (Fig. 6b; Supplementary Table 9; Supplementary Note 5). SMOOTHY-FISH successfully detected the RNA structure, with junction-probes colocalizing with the common regime signals (Fig. 6b). We also validated that the junction probe required both 10-nt sides to bind to produce any visible signal (Supplementary Fig. 10a, b). However, the junction-probe signal only colocalized with 35% of the common region, indicating that a substantial fraction of transcripts may contain alternative conformational structures (Fig. 6c; Supplementary Note 5).

We further investigated the components of the MS2-3-PP7-NiNTA structure to see which conformations the remaining transcripts could adopt. Depending on the combination of the folding and unfolding of its four modules, MS2-PP7-NiNTA can have 7 conformations, each associated with a distinct junction sequence (Fig. 6c). Conformation *I* is the expected fully folded structure, with Conformation *II* having the basal stem unfolded (Supplemental Fig. 11a, b). Conformations *III* and *IV* contain an unfolded basal stem together with an unfolded MS2 or NiNTA hairpin, respectively (Supplemental Fig. 11c, d). Conformations *V* -*VII* contain only one folded hairpin-module - MS2, PP7, or NiNTA, respectively (Fig. 6c; Supplementary Fig 11e - f).

The existence of the 6 additional conformations were tested with their respective SMOOTHY-FISH probes independently, with the common region added as a reference (Supplementary Table 9). Conformation *I* was also included in each condition to confirm that the native structure was not altered by the additional conformation probes. Conformations *II, III, IV, VI,* and *VII* had little to no signal, with each having less than a 1% abundance (Fig. 6e; Supplementary Fig. 12a - e; Supplementary Note 5). Conformation *V*, on the other hand, showed a SMOOTHY-FISH signal that colocalized tightly with the common regime at 18% abundance (Fig. 6d, 6e; Supplemental Fig. 10c, d). Signals for conformations *I* and conformations *V* did not colocalize with each other (Fig. 6d), suggesting that conformation *V* is an alternative structure for MS2-3-PP7-NiNTA. In combination with conformation *I*, the total abundance of all conformations approaches the practical benchmark of colocalization.

Despite optimizations of the structural element *in-vitro*, RNA folding in cellular environments remains less predictable than in ideal conditions^46^. The existence of conformation *V* was surprising given the stability of the overall MS2-3-PP7-NiNTA structure *in-vitro* (Supplementary Note 5). Additionally, other conformations containing only a single hairpin module, such as conformation *VI*, did not exist despite containing a stronger hairpin structure (Supplementary Fig. 12d). The concept that a designed structure can predominantly adopt a single alternative conformation in cells rather than the intended fold or a broad ensemble is not something that in-vitro characterization had revealed (Supplementary Note 5). This demonstrates that SMOOTHY-FISH can not only detect the existence of a structure but also distinguish among conformational states of an RNA in-situ.

## Discussion

We developed SMOOTHY-FISH as an effective single-probe method for mapping short RNA motifs previously inaccessible in their native cellular and tissue contexts. Through co-optimizing modular multi-fluorophore probe design, hydrogel-based sample preparation, and flanking oligonucleotide-assisted accessibility, SMOOTHY-FISH achieved detection yields comparable to the practical benchmark of hybridization-based single-molecule imaging — the performance of a full tiled smFISH probe set. Notably, the breadth of our approach follows a single design principle rather than stacking multiple specialized techniques. Because target recognition is separated from signal generation, the same optimized signal module can be directed at any short motif simply by changing the 20-nt target-binding region. So, diverse classes of RNA features, such as isoforms, small RNAs, and RNA structural conformations, can be read out by the same single-probe architecture, instead of by multiple bespoke assays. This capability was demonstrated by the spatial mapping of three classes of RNA features: splice-isoform junctions differing by as few as 2-nt, individual microRNAs shorter than a conventional probe-set target, and the conformational junctions formed by RNA folding. In short, SMOOTHY-FISH turns short RNA motifs from a blind spot into a directly measurable feature in spatial imaging.

Beyond the current implementation, SMOOTHY-FISH could be adapted for greater flexibility, multiplexing capacity, and throughput. Rather than synthesizing a distinct probe and directly hybridizing it to each target motif, a target-specific initiator oligonucleotide could be used to bridge the RNA target and a universal readout probe (Supplementary Fig. 13a)^4–6^. This design allows different motifs to be detected using the same universal SMOOTHY-FISH probe, reducing the cost and effort associated with probe synthesis. Our preliminary trial targeting the P26 site of SDHA mRNA by an initiator and a universal readout probe showed a colocalization rate comparable to direct probe hybridization(Supplementary Fig. 13b - d, Supplementary Note 6), supporting the feasibility of this design. In future implementations, incorporating barcode sequences into initiators and combining them with sequential imaging could potentially turn SMOOTHY-FISH into a multiplexable, high-throughput platform to simultaneously map large panels of RNA molecules, such as the profile of miRNAome, splice isoform repertoire, and RNA structurome (Supplementary Note 7).

Looking forward, SMOOTHY-FISH could be adapted to map many other short motifs that remain difficult to access in situ, including PIWI-interacting RNAs, snoRNAs, disease-associated repeat motifs, and antisense oligonucleotides^47–50^. Advances in RNA profiling resolution have repeatedly reshaped our understanding of RNA biology, from bulk transcript abundance to single-molecule and spatially resolved gene expression. SMOOTHY-FISH adds a further axis of measurement by extending spatial RNA profiling from long transcripts detection to short motif identification and quantification. Setting up a framework for studying RNA at a finer molecular scale, we foresee a wide range of applications that can expand the scope of spatial transcriptomics and transform our understanding of RNA biology.

## Supporting information

Manuscript with figures

## Methods and Materials

### Cell lines

HEK293 cells were obtained from ATCC (CRL-1573). HEK293T cells expressing the MS2-3-PP7-NiNTA RNA structural element were generously provided by the Prescher Lab and Lupták Lab at the University of California, Irvine (for more details regarding this cell line, see45). SH-SY5Y cells were generously provided by the Thompson Lab at the University of California, Irvine. HeLa cells were generously provided by the Shi Lab at the University of California, Irvine.

### SMOOTHY-FISH probe design and synthesis

Probe sequences used for short RNA transfection and mapping in Fig. 2f, for flanker and yield measurement in Fig. 3, for PKM1/2 isoform mapping in Fig. 4, for miRNA mapping in Fig. 5, for MS2-3-PP7-NiNTA conformation mapping in Fig. 6 are listed in Supplementary Table 4, 5 and 6, 7, 8, and 9, respectively. SMOOTHY-FISH probes were synthesized by primer extension using Deep Vent® (exo-) DNA Polymerase (NEB, M0259L). Briefly, the 25-µL reaction consisted of 4 µM template oligonucleotide, 4 µM target primer, 1× ThermoPol buffer, 400 µM each dATP, dCTP, and dGTP, 100 µM fluorophore-conjugated dUTP, and 2 units of Deep Vent® (exo-) DNA Polymerase. The primer extension reaction was performed using the following thermocycling program: initial denaturation at 95°C for 3 min; 45 cycles of annealing at 57°C for 30 s and extension at 72°C for 5 min; final extension at 72°C for 5 min; and hold at 4°C. The synthesized SMOOTHY-FISH probes were separated by gel electrophoresis and excised at the expected product size. Gel-extracted probes were purified using the QIAquick Gel Extraction Kit (QIAGEN, 28704) and eluted in DEPC-treated water. Probe concentrations were measured using a NanoDrop spectrophotometer.

### SMOOTHY-FISH protocol

#### Surface treatment of glass-bottom plates

1.5H glass-bottom 12-well and 24-well plates (Cellvis, P12-1.5H-N and P24-1.5H-N) were treated with Bind-Silane (SigmaAldrich, 440159) to introduce methacrylate groups onto the glass surface and promote hydrogel attachment. Wells were briefly rinsed with 100% ethanol, incubated with 1% Bind-Silane in 80% ethanol for 30 min at room temperature, and then rinsed twice with 100% ethanol. Plates were air-dried and baked at 95°C for 30 min on a hot plate. After cooling, wells were rinsed once with DPBS and coated with 0.1 mg/mL poly-D-lysine and 2 µg/mL human fibronectin in DPBS for 1 h at 37°C to promote cell attachment except for HeLa cells. After coating, wells were rinsed twice with 1× DPBS. Surface-treated plates were prepared immediately before use.

#### Cell culture

HEK293 cells, HeLa cells, 3T3 cells, and HEK293T cells expressing MS2-3-PP7-NiNTA were cultured in DMEM supplemented with 10% fetal bovine serum and 1% penicillin-streptomycin. SH-SY5Y cells were cultured in 1:1 DMEM/F12 supplemented with 10% fetal bovine serum and 1% penicillin-streptomycin. For SMOOTHY-FISH experiments, cells were seeded onto surface-treated glass-bottom 24-well plates at a density of 4 × 10⁴ cells per well and cultured for 16 h before fixation.

#### Fixation and permeabilization

Cell culture medium was removed, and cells were rinsed once with DPBS. Cells were fixed with 4% formaldehyde for 10 min at room temperature and washed three times with DPBS. Cells were then permeabilized with 0.5% Triton X-100 in DPBS for 10 min at room temperature and washed once with DPBS. Samples were then processed directly for in situ poly(A) tailing in miRNA experiments or for RNA poly(dT) anchoring in other SMOOTHY-FISH experiments.

#### In situ poly(A) tailing, RNA poly(dT) anchoring, and flanking oligonucleotide hybridization

For miRNA experiments, fixed and permeabilized cells were first equilibrated with E. coli poly(A) tailing buffer. Each well was then incubated with 500 µL of poly(A) tailing reaction solution containing 1× E. coli poly(A) tailing buffer, 1 mM ATP, 0.4 U/µL murine RNase inhibitor (NEB, M0314L), and 0.1 U/µL E. coli poly(A) polymerase (NEB, M0276L) at 37°C for ≥16 h. After poly(A) tailing, the reaction solution was removed, and cells were washed with 2× SSC for 15 min at room temperature on a benchtop shaker, followed by washing with wash buffer containing 2× SSC and 20% (v/v) formamide.

For RNA anchoring and flanking oligonucleotide hybridization, 3.3 µM acrydite-modified poly(dT) anchor and 20 nM each flanking oligonucleotide, when applicable, were added to each well in hybridization buffer containing 2× SSC, 20% (v/v) formamide, 1% (w/v) dextran sulfate sodium, 1 mg/mL yeast tRNA (Thermo Fisher, 15401029), 2 mM ribonucleoside vanadyl complex (NEB, S1402S), 0.1% Tween-20, and 0.4 U/µl murine RNase inhibitor. Samples were incubated at 30°C for ≥16 h.

#### Hydrogel embedding and digestion

After RNA anchoring, the anchoring solution was removed, and samples were rinsed once with wash buffer. Samples were then washed with wash buffer at 30°C for 30 min, rinsed once with 2× SSC, and kept in 2× SSC until hydrogel embedding.

The pre-gelation solution was prepared on ice by combining 7.8 mL DEPC-treated water, 0.6 mL 5 M NaCl, 0.6 mL 1 M Tris-HCl pH 8.0, 1 mL 40% 19:1 acrylamide/bis-acrylamide solution, and 2.5 µL murine RNase inhibitor. The pre-gelation solution was degassed under vacuum for 10 min and kept on ice. A fresh 10% ammonium persulfate solution was prepared and kept on ice. Circular 12-mm coverslips were treated with Gel-Slick (Lonza, 50640) to prevent adhesion to the hydrogel after polymerization.

For hydrogel embedding, plates were placed on ice, and 2× SSC was removed from the wells. Pre-gelation solution was added to each well and incubated for 5 min. Immediately before polymerization, 6 µL TEMED and 60 µL 10% ammonium persulfate were added to 10 mL pre-gelation solution and mixed to generate the gelation solution. The pre-gelation solution was removed from each well, and wells were briefly rinsed with gelation solution. Gelation solution was then added to each well, and a Gel-Slick-treated coverslip was placed into the well to contact the glass-bottom surface. Excess gelation solution was removed, leaving a thin layer of gelation solution between the coverslip and the bottom of the well. Plates were transferred to a nitrogen chamber, purged twice with nitrogen gas, and incubated at room temperature for 3 h to allow hydrogel polymerization.

After polymerization, 2× SSC was added to each well and incubated for 3 min. The 2× SSC was removed, and coverslips were carefully detached using a syringe needle and tweezers. Samples were incubated with 1 mL digestion buffer containing 2× SSC, 0.5% Triton X-100, and 1% SDS for 5 min. The buffer was then replaced with 1 mL digestion buffer supplemented with 16 U/mL proteinase K (NEB, P8107S), and samples were digested at 37°C for ≥18 h.

#### Probe hybridization

After digestion, the digestion buffer was removed, and samples were rinsed once with 2× SSC. Samples were then washed four times with 2× SSC for 15 min each at room temperature on a benchtop shaker. After the final wash, samples were equilibrated in wash buffer for 5 min. Hybridization buffer containing 0.4 U/µl murine RNase inhibitor, 20 nM each SMOOTHY-FISH probe, and/or 1 ng/µL total smFISH probe set was added to each well. Samples were incubated at 30°C for ≥18 h.

#### Post-hybridization washes

After hybridization, the hybridization buffer was removed, and samples were rinsed once with wash buffer. Samples were then washed in wash buffer at 30°C for 30 min. The wash buffer was removed, and samples were rinsed once with 2× SSC, followed by two washes with 2× SSC for 30 min each at room temperature on a benchtop shaker. Samples were stained with 1 µg/mL DAPI in 2× SSC for 30 min, rinsed once with 2× SSC, and kept in 2× SSC during imaging.

### Mouse brain tissue sample processing

Mouse brain tissue samples were generously provided by the Xu Lab at the University of California, Irvine. Briefly, a 3-month old C57/BL6 male mouse was euthanized and perfused with PBS followed by 4% PFA in 1 x PBS. Mouse brains were then harvested and then further fixed with 4% PFA overnight at 4℃. Mouse brains were then soaked in 20% sucrose for 6 hours before being embedded by OCT and flash frozen. Coronal brain sections were prepared using a Leica CM1850 cryostat at 20 µm thickness and stored in 70% ethanol at -20℃.

Sections were then transferred to 2 ml microcentrifuge tubes before undergoing permeabilization, poly(A) tailing, poly(dT) anchoring, and flanking oligonucleotide hybridization as described in cells. Tissue sections were then rinsed once with wash buffer, washed twice with wash buffer for 30 minutes at 30℃, rinsed once with 2X SSC, and washed with 2X SSC for 10 minutes. Pre-gelation solution was made and 18-mm coverslips were coated as described in cells. Tissue sections were then rinsed with the pre-gelation buffer for 30 minutes on ice before being transferred to the coated 12-well plate to air dry. Gelation was then completed as described in cells.

After polymerization, coverslips were removed and samples were rinsed with 2× SSC for 10 min. Samples were incubated in a digestion buffer containing 2× SSC, 0.5% Triton X-100, and 2% SDS for 10 min, followed by digestion with 16 U/mL proteinase K in the same buffer for two consecutive rounds (6 h and 18 h at 37°C), with a buffer rinse between rounds. Samples were then transferred to a second digestion buffer containing 2× SSC, 0.5% Triton X-100, and 0.5% SDS and digested with 32 U/mL proteinase K for two consecutive rounds (6 h and 18 h at 37°C), with a buffer rinse between rounds.

Subsequent probe hybridization and post hybridization washes were performed as described in cells.

### Image acquisition

Images were acquired in widefield fluorescence mode using an Olympus IX83 inverted microscope equipped with a 60× oil-immersion objective, an X-Cite Xylis LED illumination source (Excelitas Technologies), a Hamamatsu C13440-20CU digital camera, and MetaMorph Advanced Acquisition software. Images were collected as z-stacks with a step size of 0.4 µm. DAPI images were acquired using 10% excitation intensity and 20-ms exposure time, whereas all other fluorescence channels were acquired using 100% excitation intensity and 200-ms exposure time. Imaging settings were kept constant within each experiment. Z-stacks were converted to two-dimensional images using maximum-intensity z-projection for downstream analysis.

### In vitro poly(A) tailing and PAGE analysis

C. elegans miR-54-3p mimic RNA oligonucleotide (100 pmol) was subjected to poly(A) tailing in a 20-µL reaction for 16 h at 37°C. The reaction contained 1× E. coli poly(A) tailing buffer, 1 mM ATP, 0.4 U/µL murine RNase inhibitor, and 0.1 U/µL E. coli poly(A) polymerase. Poly(A)-tailed products were separated on a 15% denaturing TBE-urea PAGE gel (Bio-Rad, 4566053). The gel was stained with 1× SYBR™ Gold Nucleic Acid Gel Stain (Thermo Fisher, S11494) and imaged using a ChemiDoc Imaging System (Bio-Rad) with the SYBR Gold filter. Gel images were analyzed using Image Lab software (Bio-Rad).

### Transfection of miRNA mimics into HKE293 cells for SMOOTHY-FISH mapping

HEK293 cells were seeded onto silanized and coated glass-bottom 24-well plates and cultured for 16 h before transfection. Cells were transfected with 0–5 nM C. elegans miR-54-3p mimic RNA oligonucleotide using Lipofectamine™ RNAiMAX (Thermo Fisher, 13778075) according to the manufacturer’s protocol. Cells were incubated with the medium containing the transfection mixture for 2 h, fixed, and processed as described in the SMOOTHY-FISH protocol.

### RT-PCR and restriction digestion analysis of PKM1/2 isoform ratio

Total RNA was extracted from HEK293 and SH-SY5Y cells using the RNeasy Mini Kit (QIAGEN, 74104) with on-column genomic DNA removal. For tissues, RNA was extracted using the RNeasy FFPE Kit (QIAGEN, 73504) by the UCI iLAB facility. cDNA was synthesized from 1 µg total RNA using the iScript™ cDNA Synthesis Kit (Bio-Rad, 1708890) according to the manufacturer’s protocol. PCR was performed using one-tenth of the cDNA reaction in a 50-µL reaction containing 1× Q5 Reaction Buffer, 0.2 mM each dNTP, 0.5 µM each forward and reverse primer, and 1 unit of Q5® High-Fidelity DNA Polymerase. PCR amplification was performed using the following thermocycling program: initial denaturation at 98°C for 1 min; 35 cycles of denaturation at 98°C for 10 s, annealing at 63°C for 30 s, and extension at 72°C for 1 min; final extension at 72°C for 2 min; and hold at 4°C. PCR products were purified using the QIAquick PCR Purification Kit (QIAGEN, 28104). Purified PCR products (400 ng) were digested at 37°C for 1 h, using PsiI-v2 (NEB, R0744S) or PstI-HF (NEB, R3140S) for human cell line PKM analysis; using PsiI-v2 (NEB, R0744S) or SfoI (NEB, R0606S) for mouse brain tissue PKM analysis. Digested products were separated on a 1% agarose gel, stained with 1× ethidium bromide (Thermo Fisher, 15585011), and imaged using a ChemiDoc Imaging System (Bio-Rad). PKM1:PKM2 ratios were calculated from band intensities using Image Lab software (Bio-Rad), with band intensities corrected for fragment size.

### Hybridization and washing conditions for MS2-3-PP7-NiNTA mapping

For MS2-3-PP7-NiNTA conformational mapping, wash and hybridization conditions were modified from the standard SMOOTHY-FISH protocol. The formamide concentration was increased to 30% (v/v) in both wash buffer and hybridization buffer, and SMOOTHY-FISH probes were hybridized at 37°C to maintain high hybridization stringency for the GC-rich target sequences. In addition, 1 mM MgCl₂ was included in all wash and hybridization steps to help maintain RNA structure during processing.

### smFISH probe design and coupling

smFISH probes targeting SDHA mRNA were designed using Stellaris Probe Designer v4.1. Probes for human SDHA were 20-nt in length and were ordered with a 5′-AmMC6 modification. Probes for mouse SDHA were 20-nt in length and were ordered with a 3’-AmMO modification.

Amino-modified probes were coupled to fluorophore NHS esters (Lumiprobe) and purified by HPLC. Detailed probe sequences are listed in Supplementary Table 1, 2, and 6.

## Acknowledgements

We thank the Prescher and Lupták laboratories at the University of California, Irvine, for providing HEK293T cells expressing the MS2-3-PP7-NiNTA RNA structural element, the Thompson laboratory for providing SH-SY5Y cells, and the Shi laboratory for providing HeLa cells. We thank Xiaoyu Shi, John Chaput, Noah Setterholm, Stacy Copp, Chang Liu, Yongsheng Shi, Andrej Lupták, Jennifer A. Presecher, and Ding lab members for helpful discussions and suggestions during the development of SMOOTHY-FISH. We also acknowledge the Genomics Research and Technology Hub of UCI iLabs for assistance with RNA extraction and purification of tissue samples.

## Funding Statement

This work was supported by: A.D. (NIH National Eye Institute T32 EY032448, UCI VSTP Training Grant), F.D. (NIH Director’s New Innovator Award DP2GM149554, NSF DMS/NIGMS 2451263, Hypothesis Fund).

## Author Contributions

G.L. and F.D. conceived the idea and designed experiments. G.L. performed all the cellular experiments, and associated data analysis. K.J. performed all the tissue experiments, and associated data analysis. K.S. and V.K. performed the experiments for brightness quantification. G.L., K.S., and V.K. performed data analysis for brightness quantification. A.D. performed experiments on the initiator design, and associated data analysis. X.X. and Z.T. prepared all tissue samples used for the experiments. G.L. and K.J contributed equally to writing. F.D. supervised all aspects of the project.

## Competing Interest Declaration

We have patent applications (UC-2026-682-1) on the work described here.

## Additional Information

Correspondence and requests for materials should be addressed to G.L., K.J. or F.D.

