## Supplementary material for "Single-probe, high-yield spatial mapping of diverse short RNA motifs": Manuscript with figures

### Figure Legends:

**Figure 1. A single-probe strategy—SMOOTHY-FISH—expanded the scope of single-molecule spatial RNA mapping.**

**a**, The scope of single-molecule spatial RNA mapping is constrained by both the RNA molecular length and the target-specific motif length. Current tools relying on multiple probes cover only long motifs residing in long RNAs (mRNAs) while our single-probe strategy — SMOOTHY-FISH — extends the coverage to short motifs in both short RNAs (small RNAs) and long RNAs with short motifs (RNA structures, RNA isoforms). **b**, A reliable single-probe strategy needs to satisfy three requirements: probe design to enhance signal brightness, binding specificity to reduce non-specific or off-targeting signals, and target accessibility optimization to maximize detection yield. **c**, SMOOTHY-FISH's detection yield approached the practical ceiling set by the smFISH benchmark. ① SMOOTHY-FISH's detection yield was measured through colocalization between SMOOTHY-FISH (green) and smFISH (red) on SDHA mRNAs. The benchmarks were measured through colocalization between two sets of smFISH on SDHA mRNAs: ② upstream (green) and downstream (red) probe set; ③ odd (green) and even (red) probe set. Sequences of SMOOTHY-FISH and smFISH probes are listed in Supplementary Table 1, 2, & 5. In the image of SDHA mapping in an HEK293 cell in ①, fluorescent signals corresponding to smFISH (red) and SMOOTHY-FISH (green) are shown in zoom insets, with colocalized dots circled in orange and non-colocalized dots circled in blue. Detection yields in ①, ②, ③ are shown as mean ± SD (50 cells). The benchmark is shown as mean (dash line) ± SD (gray zone). See more on detection yield in Supplementary Note 1.

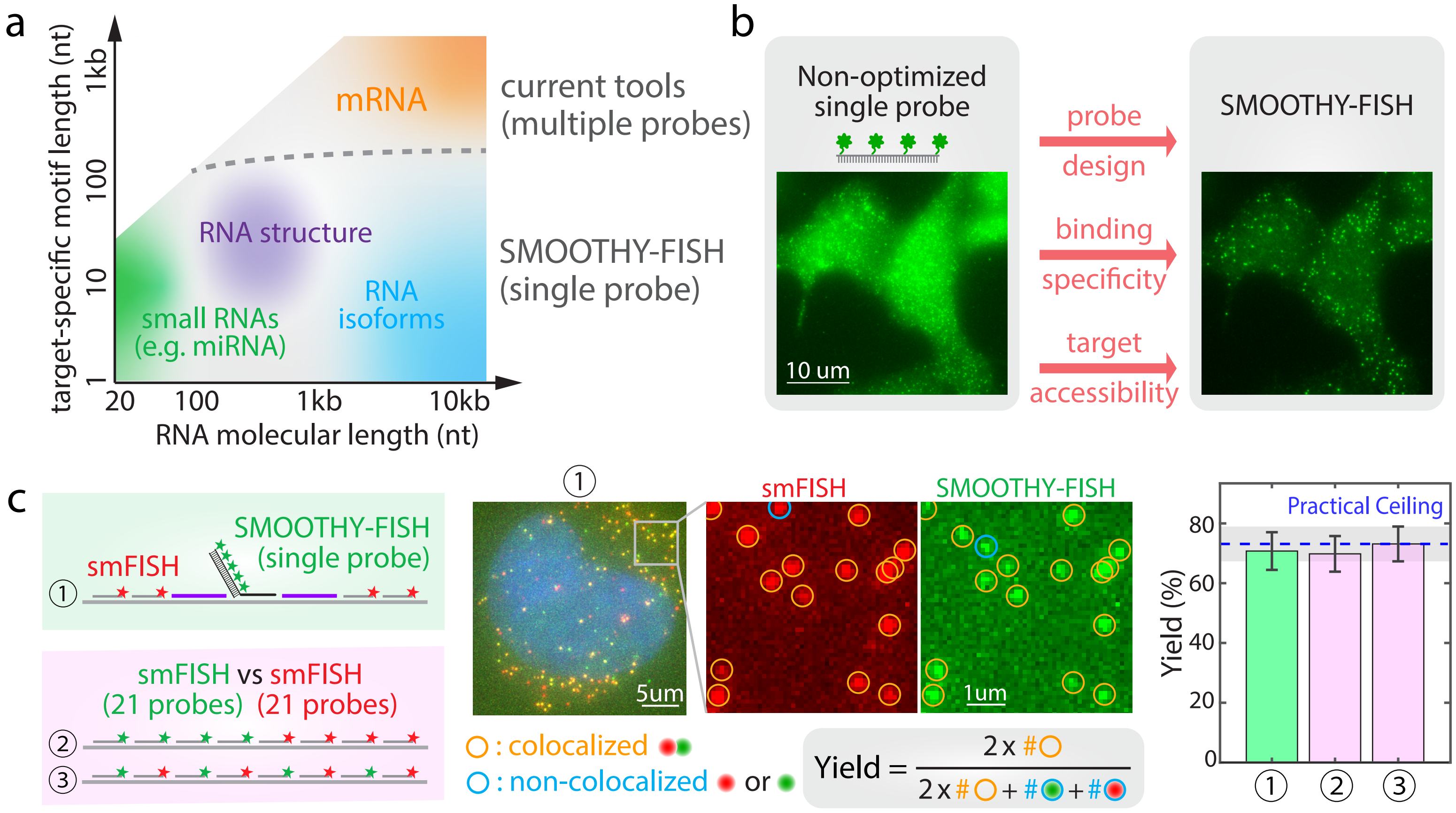
a
target-specific motif length (nt)
1kb
100
10
1
20
100
1kb
10kb
RNA molecular length (nt)
mRNA
RNA structure
small RNAs
(e.g. miRNA)
RNA
isoforms
current tools
(multiple probes)
SMOOTHY-FISH
(single probe)
b
Non-optimized
single probe
10 um
probe
design
binding
specificity
target
accessibility
SMOOTHY-FISH
c
smFISH
SMOOTHY-FISH
(single probe)
smFISH vs smFISH
(21 probes) (21 probes)
5um
smFISH
SMOOTHY-FISH
1um
: colocalized
: non-colocalized
or
Yield =
2 x #
2 x # + # + #
Practical Ceiling
Yield (%)
80
60
40
20
0

**Figure 2. Optimization of SMOOTHY-FISH probe design and binding specificity.**

**a**, The SMOOTHY-FISH probe design consists of three modules: a ~ 20-nt single-stranded target-binding region, a double-stranded signal generation region bearing multiple fluorophores, and a flexible linker to avoid steric hindrance in hybridization. **b**, SMOOTHY-FISH probes were optimized by altering the combination of fluorophore number (n) and the gap size (nt) between neighboring fluorophores. The optimal combination is represented in red in the graph. Images of suboptimal probe designs can be found in Supplementary Fig. 1. Brightness quantification can be found in Supplementary Table 3, Supplementary Note 1 and 2. **c**, Detection yield comparison among 5-nt gap probes showed 5 fluorophores with 5-nt gap achieved the highest detection yield. Detection yields are represented by mean ± SD (n = 50 cells). Notably, detection yield dropped drastically with 8 or more fluorophores on the probes. **d**, The optimal design (n = 5; gap = 5nts) was compatible with various fluorophores. We mapped SDHA mRNA by a single SMOOTHY-FISH probe with Alexa Fluor 488 (green), Alexa Fluor 594 (yellow), and ATTO647N (red). **e**, SMOOTHY-FISH's small RNAs retention in hydrogel. We verified that small RNA oligonucleotides could be poly(A)-tailed by E.coli poly(A) polymerase. PAGE left lane: negative control without E.coli poly(A) polymerase; right lane: poly(A)-tailing on a 24-nt RNA oligonucleotide. After in situ poly(A) tailing, 5'-poly(dT) oligonucleotides were applied to hybridize to poly(A) tails, followed by polyacrylamide hydrogel embedding and clearing to remove proteins and lipids. **f**, SMOOTHY-FISH signals were linearly correlated to the transfection concentration when targeting the transfected 24-nt orthogonal RNA oligonucleotides in HEK293 cells. SMOOTHY-FISH signals were represented by mean ± SD (n = 50 cells). Sequences of the orthogonal RNA oligonucleotide and the corresponding probe used in **e** and **f** are stated in Supplementary Table 4.

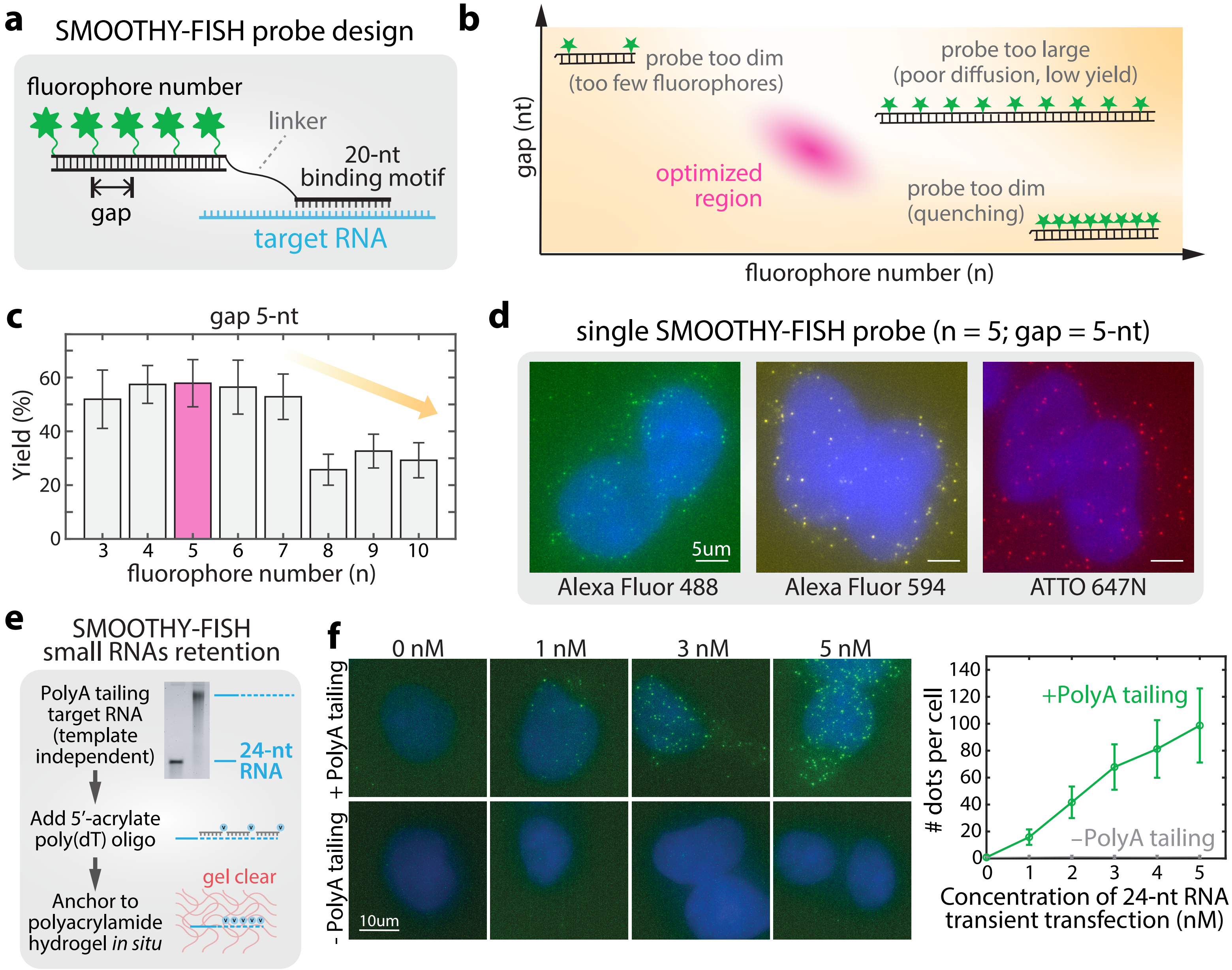
a
SMOOTHY-FISH probe design
fluorophore number
linker
20-nt
binding motif
gap
target RNA
b
probe too dim
(too few fluorophores)
probe too large
(poor diffusion, low yield)
optimized
region
probe too dim
(quenching)
gap (nt)
fluorophore number (n)
c
gap 5-nt
Yield (%)
0
20
40
60
3
4
5
6
7
8
9
10
fluorophore number (n)
d
single SMOOTHY-FISH probe (n = 5; gap = 5-nt)
5um
Alexa Fluor 488
Alexa Fluor 594
ATTO 647N
e
SMOOTHY-FISH
small RNAs retention
PolyA tailing
target RNA
(template
independent)
24-nt
RNA
Add 5′-acrylate
poly(dT) oligo
gel clear
Anchor to
polyacrylamide
hydrogel in situ
f
0 nM
1 nM
3 nM
5 nM
+ PolyA tailing
- PolyA tailing
10um
### dots per cell
0
20
40
60
80
100
120
140
+PolyA tailing
−PolyA tailing
0
1
2
3
4
5
Concentration of 24-nt RNA
transient transfection (nM)

**Figure 3. Optimization of SMOOTHY-FISH target accessibility by flanking oligonucleotides.**

**a**, We hybridized a pair of flanking oligonucleotides next to the target site to optimize the target accessibility for SMOOTHY-FISH probe hybridization. **b**, SMOOTHY-FISH mapping of site P07, P17, and P26 on SDHA mRNA showed that flankers greatly increased the detection yields. Comparison among the distance between flankers and target site showed that detection yield increase was positively correlated with the proximity in less-accessible sites (P07, P17). Detection yields were represented by mean ± SD (n = 50 cells). Sequences of smFISH probes, SMOOTHY-FISH probes, and flankers are listed in Supplementary Table 1, 2, & 5. **c**, Flankers immediately adjacent to site P17 maximized the colocalization between SMOOTHY-FISH (green) and smFISH (red) in SDHA mRNA mapping. Colocalized dots are circled in yellow; non-colocalized dots are circled in blue. **d**. SMOOTHY-FISH (green) mapping of mmuSDHA site P12 showed high colocalization to smFISH (red) in mouse brain cortex. Sequences of SMOOTHY-FISH probes, smFISH probes, and flankers are listed in Supplementary Table 6.

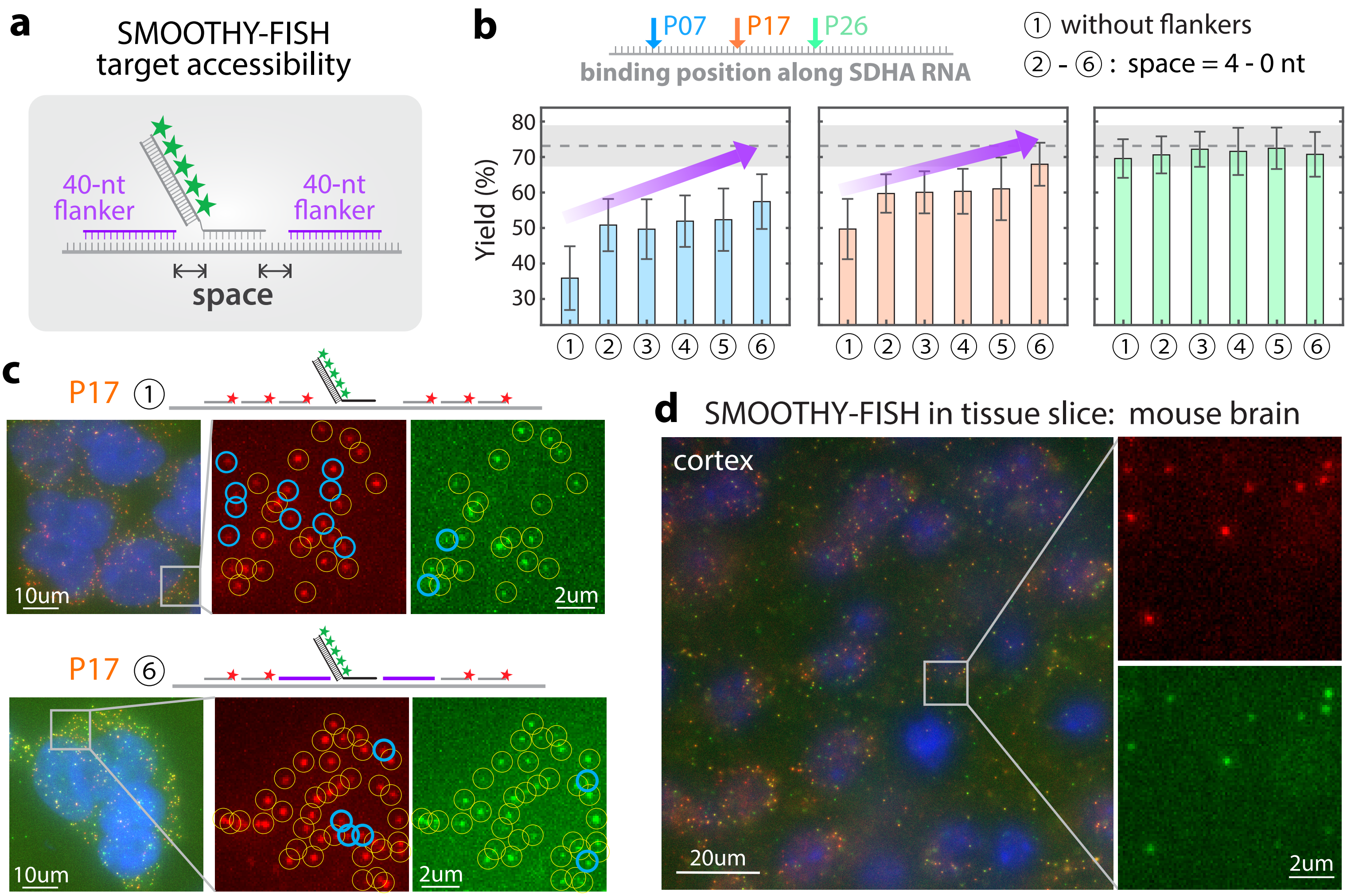
a
SMOOTHY-FISH
target accessibility
40-nt
flanker
40-nt
flanker
space
b
P07
P17
P26
binding position along SDHA RNA
① without flankers
② - ⑥ : space = 4 - 0 nt
Yield (%)
80
70
60
50
40
30
① ② ③ ④ ⑤ ⑥
c
P17 ①
10um
2um
P17 ⑥
10um
2um
d
SMOOTHY-FISH in tissue slice: mouse brain
cortex
20um
2um

**Figure 4. SMOOTHY-FISH revealed a tight expression ratio between PKM1 and PKM2 isoforms despite a wide range of expression levels in cell lines and in mouse brain tissue.**

**a**, Alternative splicing of PKM generates two isoforms PKM1 (blue, containing exon 9) and PKM2 (orange, containing exon 10) that differ by only two nucleotides. **b**, SMOOTHY-FISH probe targeted the PKM1 (green) and PKM2 (red) spliced exon junction. **c**, SMOOTHY-FISH distinguished PKM1 (green) and PKM2 (red) in HEK293 cells. Notably, PKM1 and PKM2 dot signals did not overlap, showing the high specificity of SMOOTHY-FISH. Zoomed image insets showed PKM2 was predominantly expressed in cells with different total PKM expression levels. **d**, The expression levels of PKM1 and PKM2 exhibited high variation within the cell population. PKM1 and PKM2 dot counts per cell were represented by mean ± SD in the bar graph (n = 50 cells). **e**, The ratio of Pkm1 (green) and Pkm2 (red) was flipped in the mouse hippocampus compared to cultured cell lines. Zoomed image inset showed Pkm1 was predominantly expressed across different regions in the mouse hippocampus. Sequences of SMOOTHY-FISH probes and flankers are listed in Supplementary Table 7. **f**. PKM1 and PKM2 expression ratios were very tight despite high variation in the total PKM expression across individual cells. The PKM2 fractions were estimated as the slopes of the linear fitting. The tight linear correlations are represented by the $R^2$ from the linear fitting. See the PKM2 fraction calculation in Supplementary Note 3.

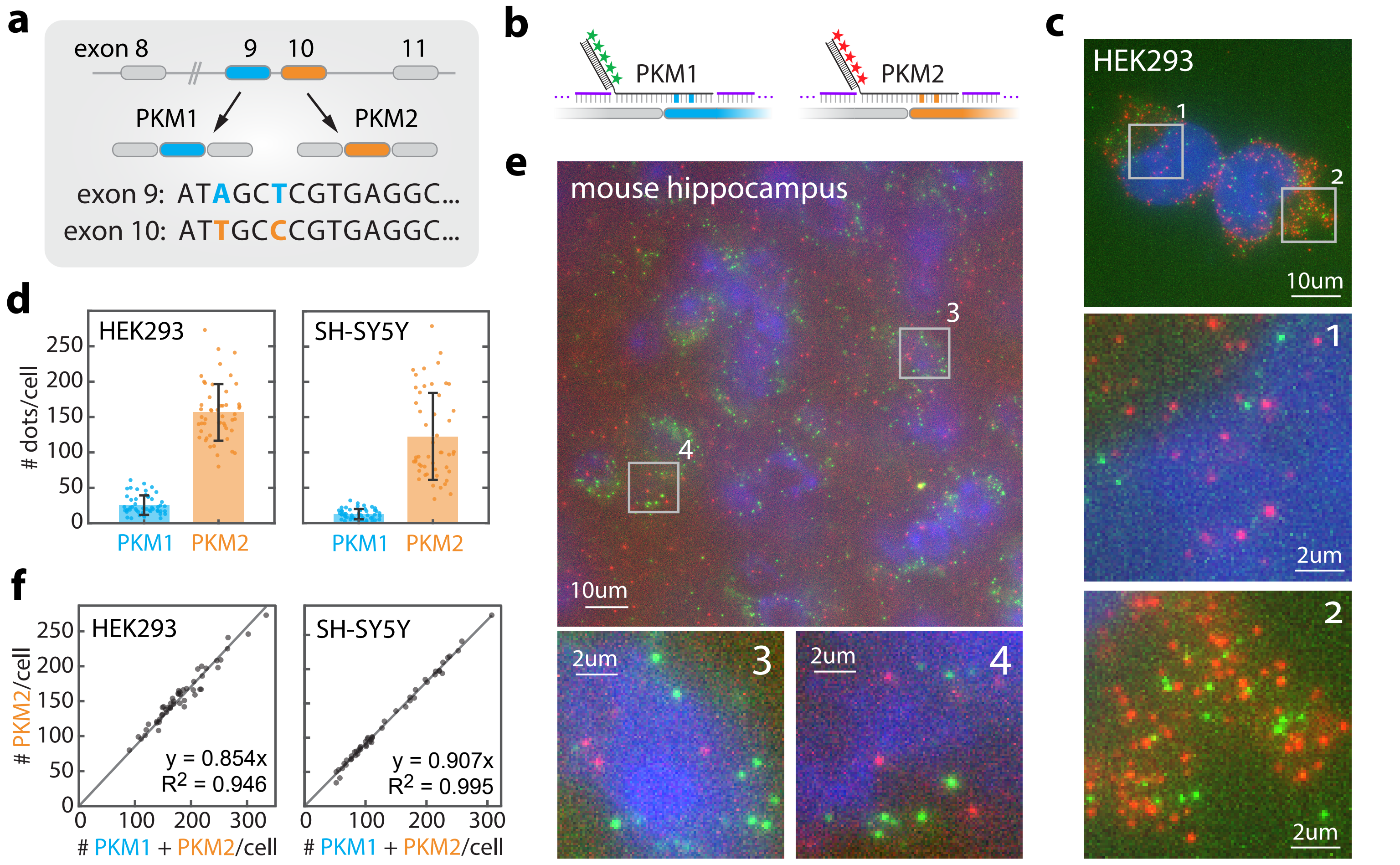
a
exon 8
9
10
11
PKM1
PKM2
exon 9: ATAGCTCGTGAGGC...
exon 10: ATTGCCCGTGAGGC...
b
PKM1
PKM2
c
HEK293
1
2
10um
2um
d
HEK293
SH-SY5Y
### dots/cell
250
200
150
100
50
0
PKM1
PKM2
e
mouse hippocampus
3
4
10um
2um
f
### PKM2/cell
### PKM1 + PKM2/cell
y = 0.854x
R2 = 0.946
y = 0.907x
R2 = 0.995

**Figure 5. SMOOTHY-FISH demonstrated that miRNAs predominantly localized to cytoplasm at tight fractions albeit the expression variation across individual cells.**

**a**, SMOOTHY-FISH was able to map miRNAs with a wide range of abundance in cells. miR-10a-5p, miR-16-5p, miR-23a-3p were reported to have low, intermediate, and high expression in HeLa cells respectively. SMOOTHY-FISH mapped these three miRNAs with and without the poly(A) tailing process. miRNA dot counts per cell are represented by mean ± SD in the bar graph (n = 50 cells). SMOOTHY-FISH mapped hippocampus-enriched miR-218-5p in **b**, hippocampus, **c**, midbrain, and **d**, hypothalamus of the mouse brain tissue with and without poly(A) tailing. **e**, miR-218-5p predominantly localized to cytoplasm in 3T3 cells (n = 50 cells). poly(A) tailing was required to retain and map miRNAs in cell lines and in mouse brain tissues. **f**, All four mapped miRNAs localized to cytoplasm at tight fractions despite the expression variation across individual cells (n = 50 cells). Linear fitting was performed between total miRNAs and cytoplasmic miRNAs for miR-10a-5p, miR-16-5p, miR-23a-3p, miR-218-5p in cell lines. The tight linear correlations are represented by the $R^2$ from the linear fitting. See the PKM2 fraction calculation in Supplementary Note 4. Sequences of SMOOTHY-FISH probes for miRNA mapping are listed in Supplementary Table 8.

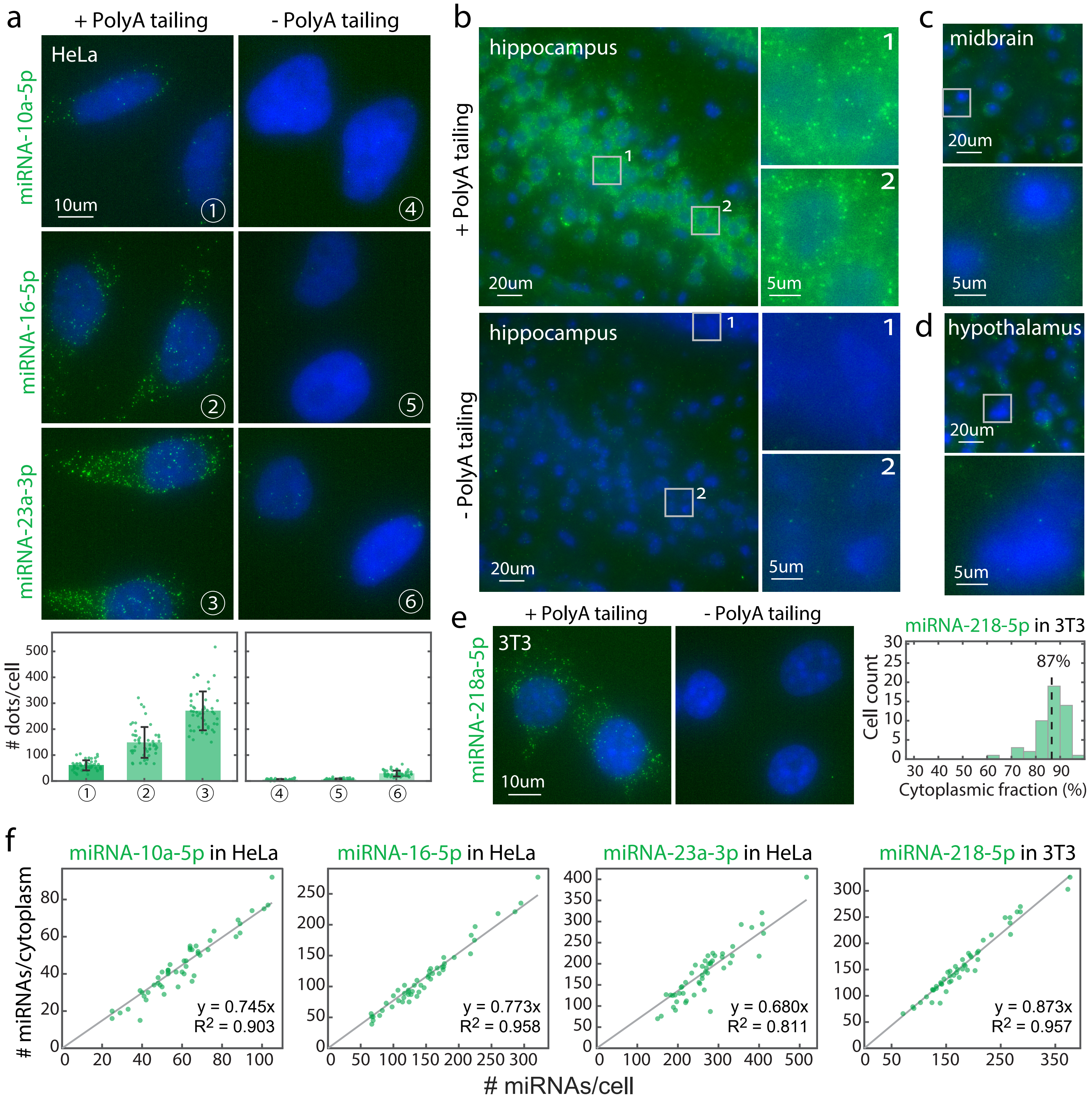

a
+ PolyA tailing
- PolyA tailing
HeLa
miRNA-10a-5p
miRNA-16-5p
miRNA-23a-3p
10um
### dots/cell
b
hippocampus
+ PolyA tailing
- PolyA tailing
20um
5um
c
midbrain
20um
5um
d
hypothalamus
20um
5um
e
+ PolyA tailing
- PolyA tailing
3T3
miRNA-218a-5p
10um
miRNA-218-5p in 3T3
87%
Cell count
Cytoplasmic fraction (%)
f
miRNA-10a-5p in HeLa
y = 0.745x
$R^2$ = 0.903
miRNA-16-5p in HeLa
y = 0.773x
$R^2$ = 0.958
miRNA-23a-3p in HeLa
y = 0.680x
$R^2$ = 0.811
miRNA-218-5p in 3T3
y = 0.873x
$R^2$ = 0.957
### miRNAs/cytoplasm
### miRNAs/cell

**Figure 6. SMOOTHY-FISH revealed an unexpected conformation of the synthetic RNA structure element MS2-3-PP7-NiNTA in its native cellular environment.**

**a**, The synthetic RNA structural element MS2-3-PP7-NiNTA, where the basal stem (orange), MS2 hairpin (pink), PP7 hairpin (yellow), and NiNTA hairpin (blue) all folded, is the most stable conformation predicted in silico (Conformation *I*). **b**, Two-colored mapping was performed by targeting a common region 5' upstream of the structural element and the Conformation *I* specific junction. SMOOTHY-FISH showed many, but not all, colocalization between the common region signals (green) and the Conformation *I* specific junction signals. Colocalized dots are circled in the zoomed image inset. The non-colocalized common region dots (green) suggested the possibility of other conformations. See more about how the junction probe sequences were designed in Supplementary Note 5. **c**, Table summary of the seven possible conformations mapped by SMOOTHY-FISH. These conformations were defined by different combinations of folded and unfolded states of the basal stem (orange), MS2 hairpin (pink), PP7 (hairpin), and NiNTA hairpin (blue). See the details of these seven conformations in Supplementary Fig. 11 and Supplementary Note 5. **d**, SMOOTHY-FISH revealed the co-existence between the Conformation *I* and Conformation *V*. Left: design of the SMOOTHY-FISH probe targeting the Conformation *V* junction (cyan). Right: three-colored SMOOTHY-FISH mapping of the common region (green), Conformation *I* junction (red), and Conformation *V* junction (cyan). Colocalized common-region and Conformation *I* dots are marked with circles and colocalized common-region and Conformation *V* dots are marked with squares in the zoomed image insets. See the three-colored SMOOTHY-FISH mapping of the common region, Conformation *I* junction, other conformation junctions in Supplementary Fig. 12. Sequences of probes for conformation mapping are listed in Supplementary Table 9. **e**, Colocalization fraction quantification of the three-colored mapping results showed Conformation *I* and Conformation *V* were the two co-existed and dominant conformations of the MS2-3-PP7-NiNTA structure element in its native cellular environment. Colocalization fractions with the common region were obtained by three-color mapping of the common region, the Conformation *I* junction, and each junction from the Conformation *II* – *VII* junctions in each experiment. Colocalization fractions are shown as medians with the $12.5^{th}$–$87.5^{th}$ percentile ranges (n = 50 cells) in the bar graph. See the calculation in Supplementary Note 5.

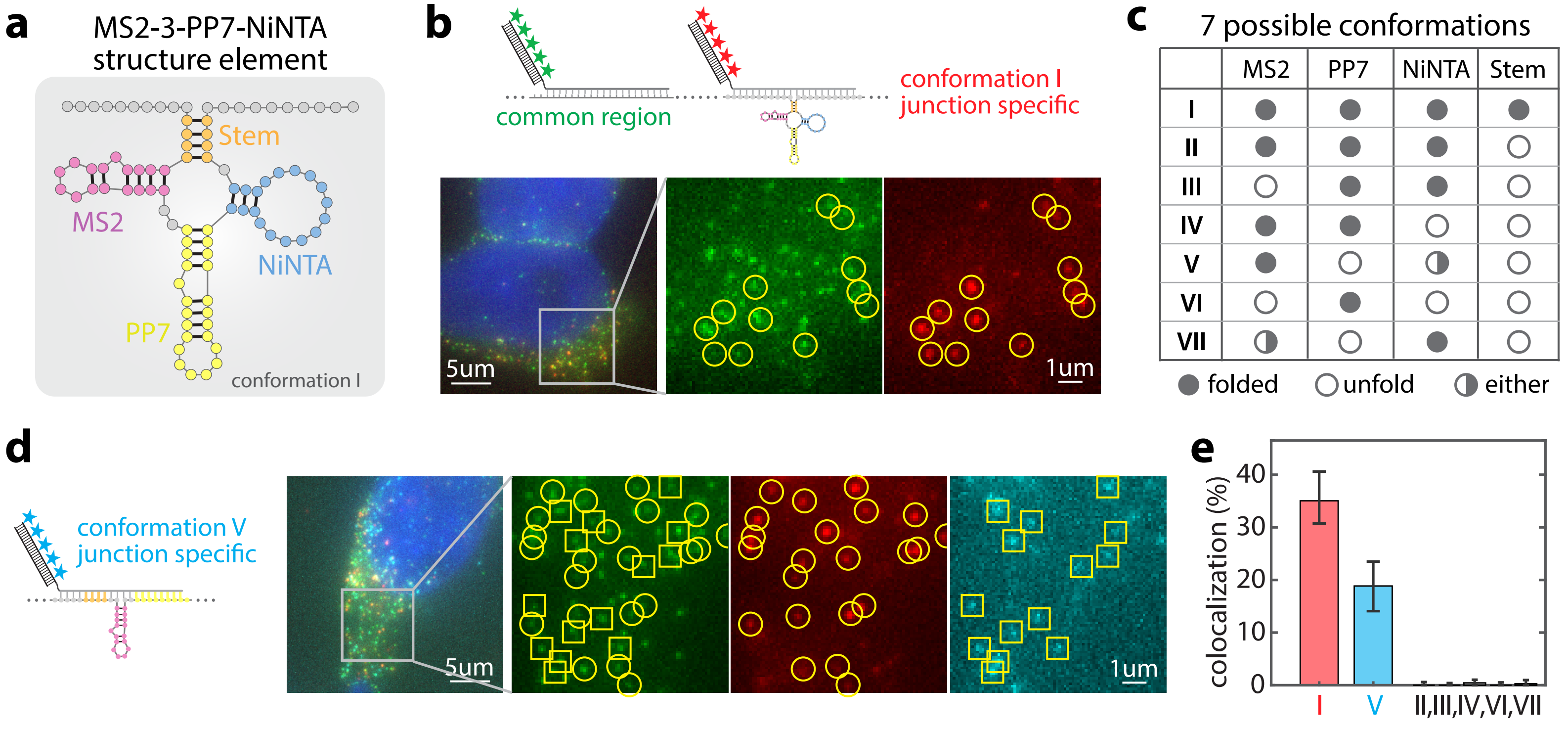
a
MS2-3-PP7-NiNTA
structure element
Stem
MS2
NiNTA
PP7
conformation I
b
common region
conformation I
junction specific
5um
1um
c
7 possible conformations
MS2
PP7
NiNTA
Stem
I
II
III
IV
V
VI
VII
folded
unfold
either
d
conformation V
junction specific
5um
1um
e
colocalization (%)
0
10
20
30
40
I
V
II,III,IV,VI,VII

### Supplementary Tables:

**Supplementary Table 1: smFISH probes targeting SDHA mRNA (Homo sapiens)**

| Oligonucleotide name | DNA oligonucleotide sequence (5'->3') |
|---|---|
| SDHA01 | /5AmMC6/AAGTGAAAACCTCGGGTTCC |
| SDHA02 | /5AmMC6/TGAAACTTTAGCAGATGCCC |
| SDHA03 | /5AmMC6/GATCCACTACTGGATACTGA |
| SDHA04 | /5AmMC6/TACCACCACTGCATCAAATT |
| SDHA05 | /5AmMC6/AGAAAGGCCAAATGCAGCTC |
| SDHA06 | /5AmMC6/CACATGCTGTATTAAACCCT |
| SDHA07 | /5AmMC6/TGTGTGACCTGGTAGGAAAC |
| SDHA08 | /5AmMC6/TCACGGTGTCGTAGAAATGC |
| SDHA09 | /5AmMC6/GTCATGTAGTGGATGGCATC |
| SDHA10 | /5AmMC6/AATTTTCTAGCTCGACCACG |
| SDHA11 | /5AmMC6/TGCACGCTGATAAATCTTCC |
| SDHA12 | /5AmMC6/TTTCCAAACTTGAGGCTCTG |
| SDHA13 | /5AmMC6/GACCTTCCATATAAGGTGTG |
| SDHA14 | /5AmMC6/ATAGCTGGTATCATATCGCA |
| SDHA15 | /5AmMC6/CATCAGGAGATCCAAGGCAA |
| SDHA16 | /5AmMC6/TATGCACAGTGCGATGACAC |
| SDHA17 | /5AmMC6/CTCTTATGCGATGGATGGAC |
| SDHA18 | /5AmMC6/TGTGGCAACAACAGTGTTCT |
| SDHA19 | /5AmMC6/CAGACGTGCAGCTGAAGTAG |
| SDHA20 | /5AmMC6/AAACTCTAGGTCCTGGCAAG |
| SDHA21 | /5AmMC6/TATATGCCTGTAGGGTGGAA |
| SDHA22 | /5AmMC6/ACATCCTTCCGTAATGAGAC |
| SDHA23 | /5AmMC6/CTGTTAATGAGAATGCCTCC |
| SDHA24 | /5AmMC6/TATCGCTCCATAAACCTTTC |
| SDHA25 | /5AmMC6/GAGACACCACATCTCTAGAC |
| SDHA26 | /5AmMC6/CAGGTAGACGTGATCTTTCT |
| SDHA27 | /5AmMC6/AGATCATGGCTGTCTCTGAA |
| SDHA28 | /5AmMC6/TTATAATGCACGGTGGGGAG |
| SDHA29 | /5AmMC6/CGACCAAAGACAACCAGGTC |
| SDHA30 | /5AmMC6/CATGACTCTTCGATGCTCAG |
| SDHA31 | /5AmMC6/GCGTTTGGTTTAATTGGAGG |
| SDHA32 | /5AmMC6/ATTCATGACAGATTCTTCCC |
| SDHA33 | /5AmMC6/CTTATGCTTCCATCAGCAAA |
| SDHA34 | /5AmMC6/ATTGACTTCTGCATGCTGAG |
| SDHA35 | /5AmMC6/CACAACCTTCTTGCAACACG |
| SDHA36 | /5AmMC6/CCATAGAGCTTGCTGATTTT |
| SDHA37 | /5AmMC6/AACGTCTTCAGGTGCTTTAG |
| SDHA38 | /5AmMC6/AATCGTACTCATCAATCCGC |
| SDHA39 | /5AmMC6/TCAAAGGGCTTCTTCTGTTG |

| | |
|---|---|
| SDHA40 | /5AmMC6/CAACGTCCACATAGGACAGG |
| SDHA41 | /5AmMC6/CTATATTCCAGAGTGACCTT |
| SDHA42 | /5AmMC6/CCTCGTTCAAAGTTTTGTCG |
| SDHA43 | /5AmMC6/TCTCATCAGTAGGAGCGAAT |
| SDHA44 | /5AmMC6/TGATTCTGTCATCACCACAT |
| SDHA45 | /5AmMC6/TGGACACATGCATGAGCTAT |
| SDHA46 | /5AmMC6/CAGAGTGCAGAAGCGTATGA |
| SDHA47 | /5AmMC6/AATCTCCCTTCAATGTACTC |
| SDHA48 | /5AmMC6/GGACAAAGGTAAGTGCCACG |

**Supplementary Table 2: SDHA smFISH probe sets (Homo Sapiens)**

| Probe set name | SDHA smFISH oligonucleotide sets |
|---|---|
| SDHA21-ATTO647N-UP | 1 - 23, skipping 7 and 17 |
| SDHA21-AF488-DOWN | 24 - 45, skipping 26 |
| SDHA21-ATTO647N-ODD | 1, 3, 5, 9, 11, 13, 15, 19, 21, 23, 25, 27, 29, 31, 33, 35, 37, 39, 41, 43, 45 |
| SDHA21-AF488-EVEN | 2, 4, 6, 8, 10, 12, 14, 16, 18, 20, 22, 24, 28, 30, 32, 34, 36, 38, 40, 42, 44 |
| SDHA-total-ATTO647N | 1 - 48, skipping 7, 17, 26, 39, 46 |

**Supplementary Table 3: Probes Brightness quantification**

| Gap size (nt) | Fluorophore | Fluorophore number (n) | Brightness median | Brightness 95% bootstrap CI |
|---|---|---|---|---|
| 5 | Alexa Fluor 488 | 3 | 352.085 | [342.259, 360.525] |
| 5 | Alexa Fluor 488 | 4 | 360.348 | [347.964, 371.966] |
| 5 | Alexa Fluor 488 | 5 | 393.175 | [382.434, 404.076] |
| 5 | Alexa Fluor 488 | 6 | 442.517 | [427.468, 452.814] |
| 5 | Alexa Fluor 488 | 7 | 473.461 | [459.474, 485.289] |
| 5 | Alexa Fluor 488 | 8 | 394.383 | [386.047, 403.138] |
| 5 | Alexa Fluor 488 | 9 | 397.201 | [388.284, 406.746] |
| 5 | Alexa Fluor 488 | 10 | 438.995 | [430.109, 448.872] |
| 5 | Alexa Fluor 594 | 3 | 775.637 | [743.784, 825.547] |
| 5 | Alexa Fluor 594 | 4 | 880.431 | [837.082, 914.254] |
| 5 | Alexa Fluor 594 | 5 | 1074.147 | [1018.742, 1122.765] |
| 5 | Alexa Fluor 594 | 6 | 906.524 | [856.013, 953.431] |
| 5 | Alexa Fluor 594 | 7 | 844.153 | [812.996, 887.069] |
| 5 | Alexa Fluor 594 | 8 | 993.897 | [963.293, 1030.116] |
| 5 | Alexa Fluor 594 | 9 | 1070.125 | [1041.364, 1107.619] |
| 5 | Alexa Fluor 594 | 10 | 1016.762 | [981.066, 1056.188] |
| 5 | ATTO647N | 3 | 436.396 | [424.064, 447.299] |
| 5 | ATTO647N | 4 | 503.277 | [493.125, 513.761] |
| 5 | ATTO647N | 5 | 795.608 | [777.377, 810.786] |
| 5 | ATTO647N | 6 | 685.685 | [671.389, 700.197] |
| 5 | ATTO647N | 7 | 744.591 | [729.209, 758.100] |
| 5 | ATTO647N | 8 | 545.215 | [536.733, 553.872] |
| 5 | ATTO647N | 9 | 790.778 | [773.674, 809.293] |
| 5 | ATTO647N | 10 | 845.812 | [826.783, 862.654] |
| 4 | Alexa Fluor 488 | 3 | 286.704 | [276.344, 294.981] |
| 4 | Alexa Fluor 488 | 4 | 360.249 | [350.668, 370.322] |
| 4 | Alexa Fluor 488 | 5 | 470.784 | [459.887, 481.259] |
| 4 | Alexa Fluor 488 | 6 | 422.892 | [410.776, 430.800] |
| 4 | Alexa Fluor 488 | 7 | 555.562 | [534.668, 572.201] |
| 4 | Alexa Fluor 488 | 8 | 732.150 | [713.188, 751.828] |
| 4 | Alexa Fluor 488 | 9 | 568.095 | [557.140, 584.463] |
| 4 | Alexa Fluor 488 | 10 | 796.120 | [773.565, 813.196] |
| 4 | Alexa Fluor 594 | 3 | 838.664 | [803.194, 872.692] |
| 4 | Alexa Fluor 594 | 4 | 766.311 | [738.373, 792.916] |
| 4 | Alexa Fluor 594 | 5 | 826.506 | [798.708, 854.600] |
| 4 | Alexa Fluor 594 | 6 | 908.231 | [877.517, 934.539] |
| 4 | Alexa Fluor 594 | 7 | 1366.778 | [1337.985, 1395.777] |
| 4 | Alexa Fluor 594 | 8 | NA[1] | NA[1] |
| 4 | Alexa Fluor 594 | 9 | NA[1] | NA[1] |
| 4 | Alexa Fluor 594 | 10 | NA[1] | NA[1] |
| 4 | ATTO647N | 3 | 338.589 | [325.073, 361.026] |
| 4 | ATTO647N | 4 | 357.332 | [348.002, 370.657] |
| 4 | ATTO647N | 5 | 347.042 | [331.698, 361.054] |

| | | | | |
|---|---|---|---|---|
| 4 | ATTO647N | 6 | 361.330 | [345.501, 379.201] |
| 4 | ATTO647N | 7 | 777.724 | [750.441, 799.843] |
| 4 | ATTO647N | 8 | 808.738 | [781.098, 839.114] |
| 4 | ATTO647N | 9 | 725.346 | [704.032, 743.682] |
| 4 | ATTO647N | 10 | 792.084 | [770.611, 811.673] |
| 3 | Alexa Fluor 488 | 5 | 329.292 | [317.299, 339.132] |
| 3 | Alexa Fluor 488 | 10 | 732.527 | [690.593, 765.509] |
| 3 | Alexa Fluor 488 | 16 | 539.833 | [517.361, 574.870] |
| 3 | Alexa Fluor 594 | 5 | 820.526 | [785.321, 851.360] |
| 3 | Alexa Fluor 594 | 10 | 1001.596 | [968.587, 1031.422] |
| 3 | Alexa Fluor 594 | 16 | 800.753 | [778.224, 819.014] |
| 3 | ATTO647N | 5 | NA[2] | NA[2] |
| 3 | ATTO647N | 10 | NA[2] | NA[2] |
| 3 | ATTO647N | 16 | NA[2] | NA[2] |

Note 1: probes aggregated and formed bright clumps in cell samples.
Note 2: probes did not generate any visible dots in cell samples.

**Supplementary Table 4: C. elegans miRNA mimic**

| **Oligonucleotide name** | **RNA oligonucleotide sequence (5'->3')** |
|---|---|
| C. elegans miR-54-3p mimic | UACCCGUAAUCUUCAUAAUCCGAG |

| **Oligonucleotide name** | **DNA oligonucleotide sequence (5'->3')** |
|---|---|
| Anti - C. elegans miR-54-3p mimic | CTCGGATTATGAAGATTACGGGTA |

**Supplementary Table 5: SMOOTHY-FISH probes targeting SDHA**

| Oligonucleotide name | DNA oligonucleotide sequence (5'->3') |
|---|---|
| P07-AF488 | TGTGTGACCTGGTAGGAAAC |
| P17-AF488 | CTCTTATGCGATGGATGGAC |
| P26-AF488 | CAGGTAGACGTGATCTTTCT |

| | Probe design | DNA oligonucleotide sequence (5'->3') |
|---|---|---|
| **flankers for P07** | Pairs for ⑥ (space = 0) | AGCTTGGTAACACATGCTGTATTAAACCCTGCCTCAGAAA |
| | | TTCCCCAGAGCAGCATTGATTCCTCCCTGTGCTGCAACAG |
| | Pairs for ⑤ (space = 1nt) | GCTTGGTAACACATGCTGTATTAAACCCTGCCTCAGAAAG |
| | | GTTCCCCAGAGCAGCATTGATTCCTCCCTGTGCTGCAACA |
| | Pairs for ④ (space = 2nts) | CTTGGTAACACATGCTGTATTAAACCCTGCCTCAGAAAGG |
| | | TGTTCCCCAGAGCAGCATTGATTCCTCCCTGTGCTGCAAC |
| | Pairs for ③ (space = 3nts) | TTGGTAACACATGCTGTATTAAACCCTGCCTCAGAAAGGC |
| | | ATGTTCCCCAGAGCAGCATTGATTCCTCCCTGTGCTGCAA |
| | Pairs for ② (space = 4nts) | TGGTAACACATGCTGTATTAAACCCTGCCTCAGAAAGGCC |
| | | CATGTTCCCCAGAGCAGCATTGATTCCTCCCTGTGCTGCA |

| | Probe design | DNA oligonucleotide sequence (5'->3') |
|---|---|---|
| **flankers for P17** | Pairs for ⑥ (space = 0) | CCGTCCTCTATGCACAGTGCGATGACACCACGGCACTCCC |
| | | GTGCGCCCGTAGCCTCCTGTGGCAACAACAGTGTTCTTTG |
| | Pairs for ⑤ (space = 1nt) | CGTCCTCTATGCACAGTGCGATGACACCACGGCACTCCCC |
| | | GGTGCGCCCGTAGCCTCCTGTGGCAACAACAGTGTTCTTT |
| | Pairs for ④ (space = 2nts) | GTCCTCTATGCACAGTGCGATGACACCACGGCACTCCCCA |
| | | AGGTGCGCCCGTAGCCTCCTGTGGCAACAACAGTGTTCTT |
| | Pairs for ③ (space = 3nts) | TCCTCTATGCACAGTGCGATGACACCACGGCACTCCCCAT |
| | | TAGGTGCGCCCGTAGCCTCCTGTGGCAACAACAGTGTTCT |
| | Pairs for ② (space = 4nts) | CCTCTATGCACAGTGCGATGACACCACGGCACTCCCCATT |
| | | GTAGGTGCGCCCGTAGCCTCCTGTGGCAACAACAGTGTTC |

| | Probe design | DNA oligonucleotide sequence (5'->3') |
|---|---|---|
| **flankers for P26** | Pairs for ⑥ (space = 0) | CAGGGCCACAGCCTCTTCCTTCTCGGATCTCCAGAGTCAT |
| | | GGCGCGTGGCCAGCTGCTCTGGAGGTAGGTGGTGCAGCTG |
| | Pairs for ⑤ (space = 1nt) | AGGGCCACAGCCTCTTCCTTCTCGGATCTCCAGAGTCATG |
| | | AGGCGCGTGGCCAGCTGCTCTGGAGGTAGGTGGTGCAGCT |
| | Pairs for ④ (space = 2nts) | GGGCCACAGCCTCTTCCTTCTCGGATCTCCAGAGTCATGG |
| | | CAGGCGCGTGGCCAGCTGCTCTGGAGGTAGGTGGTGCAGC |
| | Pairs for ③ (space = 3nts) | GGCCACAGCCTCTTCCTTCTCGGATCTCCAGAGTCATGGA |
| | | GCAGGCGCGTGGCCAGCTGCTCTGGAGGTAGGTGGTGCAG |
| | Pairs for ② (space = 4nts) | GCCACAGCCTCTTCCTTCTCGGATCTCCAGAGTCATGGAC |
| | | GGCAGGCGCGTGGCCAGCTGCTCTGGAGGTAGGTGGTGCA |

**Supplementary Table 6: Mus Musculus sequences**

| Oligonucleotide name | DNA oligonucleotide sequence (5'->3') |
|---|---|
| mmuSDHA01 | CTTGTTCTCACCAACAGAGA/3AmMO/ |
| mmuSDHA02 | TGCAACAGTATGTGATCGGG/3AmMO/ |
| mmuSDHA03 | ATAGAAATGCCATCTCCAGT/3AmMO/ |
| mmuSDHA04 | ATGTAATGGATGGCATCCTG/3AmMO/ |
| mmuSDHA05 | ATCTTCCATACAACGTGTGT/3AmMO/ |
| mmuSDHA06 | TTCCATCAGAAGATCCAGTG/3AmMO/ |
| mmuSDHA07 | GTACAGCTGAAGTAGGTTCG/3AmMO/ |
| mmuSDHA08 | TAAGTCCTGGCAAGGCAAAC/3AmMO/ |
| mmuSDHA09 | CATGAACCTTTCACCTTGAC/3AmMO/ |
| mmuSDHA10 | ACAACATCTCTTGATGCCAG/3AmMO/ |
| mmuSDHA11 | CGGATCTCAAGAGTCATGGA/3AmMO/ |
| mmuSDHA12 | CAGGTAGACGTGATCTTTCT/3AmMO/ |
| mmuSDHA13 | TCTCTGAAATTCCGGGTAGA/3AmMO/ |
| mmuSDHA14 | TTGTAATGCACAGTGGGGAG/3AmMO/ |
| mmuSDHA15 | CATACAGACCAGGCACAATC/3AmMO/ |
| mmuSDHA16 | CCAAGAGAGAATTTGCTCCA/3AmMO/ |
| mmuSDHA17 | TTCATAACCGATTCTTCTCC/3AmMO/ |
| mmuSDHA18 | CCATCGGCAAATCTCAACTT/3AmMO/ |
| mmuSDHA19 | TTAGGCGTAGTTCTGATGTT/3AmMO/ |
| mmuSDHA20 | GGGCTTGGAGTAATCATACT/3AmMO/ |
| mmuSDHA21 | GATGTCCACATATGAGAGGG/3AmMO/ |
| mmuSDHA22 | GTACTCCAAAGTAACCTTGC/3AmMO/ |
| mmuSDHA23 | AGGTCTTGTCGATAACAGGT/3AmMO/ |
| mmuSDHA24 | TTCATCAGTAGGAGCGGATA/3AmMO/ |
| mmuSDHA25 | AACGCAGGTAAGCTATGCTA/3AmMO/ |
| mmuSDHA26 | CCTTGTTCATGAGAGCAGAA/3AmMO/ |
| mmuSDHA27 | GAGACAAAGTCTGGCGCAAC/3AmMO/ |
| mmuSDHA28 | AAATTATGAAGGACCTGCCC/3AmMO/ |
| mmuSDHA29 | CTCGATACTTACCATATCCA/3AmMO/ |
| mmuSDHA30 | CCAGGAACCTAAACAAGGCA/3AmMO/ |

| Probe set name | SDHA smFISH oligonucleotide sets |
|---|---|
| mmuSDHA-total-ATTO647N | 1 - 30, skipping 2, 12, 17, 24, 29 |

| Oligonucleotide name | DNA oligonucleotide sequence (5'->3') |
|---|---|
| mmuP12-AF594 | CAGGTAGACGTGATCTTTCT |

<table>
<tr><th rowspan="3">flankers for mmuP12</th><th>Probe design</th><th>DNA oligonucleotide sequence (5'->3')</th></tr>
<tr><td rowspan="2">Pairs for ⑥ (space = 0)</td><td>CAGGGCCACAGCCTCTTCCTTCACGGATCTCAAGAGTCAT</td></tr>
<tr><td>GACGTGTGGCCAGTTGCTCAGGAGGCAGATGATGCAACTG</td></tr>
</table>

**Supplementary Table 7: Probes targeting PKM isoforms**

| Oligonucleotide name | DNA oligonucleotide sequence (5'->3') |
|---|---|
| HEK293-PKM1-AF488 | TCACGAGCTATCAGGTGCTG |
| HEK293-PKM2-AF594 | TCACGGGCAATCAGGTGCTG |
| HEK293-PKM1-AF594 | TCACGAGCTATCAGGTGCTG |
| HEK293-PKM2-AF488 | TCACGGGCAATCAGGTGCTG |
| Mouse-PKM1-ATTO647N | TCCCGAGCTATCAGGTGCTG |
| Mouse-PKM2-AF594 | TCTCGGGCAATCAGGTGCTG |
| Mouse-PKM1-AF594 | TCCCGAGCTATCAGGTGCTG |
| Mouse-PKM2-ATTO647N | TCTCGGGCAATCAGGTGCTG |

| | Probe design | DNA oligonucleotide sequence (5'->3') |
|---|---|---|
| **flankers for PKM isoforms** | Pairs for HEK293-PKM1 | CATGCGCACAGCCTCCAGAGGATAGTCCCCTTTGGCTGTT |
| | | TTCTTCAAACAGCTTGCGGTGGAACATGGCTGCCTCAGCC |
| | Pairs for HEK293-PKM2 | CATGCGCACAGCCTCCAGAGGATAGTCCCCTTTGGCTGTT |
| | | TTCCTCAAATAATTGCAAGTGGTAGATGGCAGCCTCTGCC |
| | Pairs for Mouse-PKM1 | CATGCGAACAGCCTCCAGAGGGTAGTCCCCCTTGGCTGTT |
| | | CTCTTCAAACAGCAGACGGTGGAACATGGCTGCCTCAGCC |
| | Pairs for Mouse-PKM2 | CATGCGAACAGCCTCCAGAGGGTAGTCCCCCTTGGCTGTT |
| | | TTCCTCGAATAGCTGCAAGTGGTAGATGGCAGCCTCTGCC |

| RT-PCR primer name | DNA oligonucleotide sequence (5'->3') |
|---|---|
| HEK293-PKM-F | AAGCCTGTCATCTGTGC |
| HEK293-PKM-R | GAGCTGTCTGGGGATTC |
| Mouse-PKM-F | TGTCTGGAGAAACAGCCAAG |
| Mouse-PKM-R | CATCTTCCCTGTGCTGTGTAA |

**Supplementary Table 8: Probes targeting miRNAs**

| Oligonucleotide name | DNA oligonucleotide sequence (5'->3') |
|---|---|
| mmu-miR-218-5p | ACATGGTTAGATCAAGCACAA |
| HeLa-miR-23a-3p-AF488 | TTGGAAATCCCTGGCAATGTGAT |
| HeLa-miR-16a-5p-AF488 | CGCCAATATTTACGTGCTGCTA |
| HeLa-miR-10a-5p-AF488 | CACAAATTCGGATCTACAGGGTA |

**Supplementary Table 9: Probes targeting M-3-P structures**

| Oligonucleotide name | DNA oligonucleotide sequence (5'->3') |
|---|---|
| Common region-AF488 | CGACGTAGGTCTCGTTGTTG |
| Conformation-1- AF594 | GCATGCTCGAGCGGCCGCCA |
| Conformation-2-ATTO647N | CTCGAGGGAGTCCCGCGGCC |
| Conformation-3-ATTO647N | CTCGAGGGAGTCGACGGCTG |
| Conformation-4-ATTO647N | TTCAATTGCCTCCCGCGGCC |
| Conformation-5-ATTO647N | GTCTGCTCTCTCCCGCGGCC |
| Conformation-6-ATTO647N | TTCAATTGCCTCGACGGCTG |
| Conformation-7-ATTO647N | CTCGAGGGAGGAGCGACGCC |

| | Probe design | DNA oligonucleotide sequence (5'->3') |
|---|---|---|
| **flankers for M-3-P structure** | Pairs for common region | GCCTCCTTGATTCTTTCCAGTCTGTAGTCCACATAGTAGACGCCA |
| | | TGCTAGGGAGGTCGCAGTATCTGGCCACTGCCACCTCGTGCTGCT |
| | Paris for structure region | GTGTGATGGATATCTGCAGAATTCTTAATTAAGCTTGTGCCCCAG |
| | | CAAATTTTGTAATCCAGAGGTTGATTGTCGACTTACCCTCTAGAT |

| Half junction probe oligonucleotide name | DNA oligonucleotide sequence (5'->3') |
|---|---|
| Conformation-1-upstream-ATTO647N | GCGGCCGCCA |
| Conformation-1-downstream-ATTO647N | GCATGCTCGA |
| Conformation-5-upstream-ATTO647N | TCCCGCGGCC |
| Conformation-5-downstream-ATTO647N | GTCTGCTCTC |

### Supplementary Notes:

**Supplementary Note 1: Calculation of detection yield**

**Image acquisition**

We used epifluorescence microscopy to image samples in Z-stacks with a step size of 0.4μm in 60X magnification. This step size is small enough to capture smFISH and SMOOTHY-FISH dot signals. We removed the out-of-focus slices that focused on the bottom of the well, because the positively charged poly-D-lysine coated surface harbors some non-specific binding of smFISH and SMOOTHY-FISH probes. Then, we used the Z-maximum projection to re-construct the 2D sample for RNA localization and quantification analysis. The microscopy images have the size of 2048 x 2048 pixels; each pixel length equals 108nm.

We used a MATLAB pipeline developed by Ding, F & Elowitz, M (2019) in the previous study[23] for cell segmentation, smFISH and SMOOTHY-FISH dot detection, colocalization dot pairing, and dot intensity fitting.

**Segmentation**

We manually segmented single cells on the MATLAB pipeline panel. Because our SMOOTHY-FISH workflow has a step of clearing that removes proteins and lipids, the sample autofluorescence was much lower compared to uncleared samples. For samples with high cellular autofluorescence (e.g. HEK293 cells and SH-SY5Y cells), we segmented the cell bodies according to the autofluorescence. For samples with low cellular autofluorescence (e.g. HeLa cells and 3T3 cells), we add SDHA smFISH probes during the hybridization and use the SDHA smFISH dots to guide the cell body segmentation. We segmented the nuclei by the DAPI signals. We envision that it is possible to apply functionalized membrane anchors and stains51 to further facilitate the segmentation process if samples have complex morphologies that are incompatible with autofluorescence- or FISH-dot-guided segmentation.

For tissue image analysis, singular cells could not be segmented due to the 3D architecture and dense cellular areas. Instead, whole tissue areas were segmented to mimic the capture of multiple full cells across an entire area. This was accomplished by drawing boundaries near but not through cellular nuclei to minimize cutting through different cells. For the hippocampus cornu ammonia regions, only areas with high cellular density were included to ensure that whole tissue regions contained similar cells with similar boundaries. Other brain regions had cellular compositions that were more uniform, and included as many cells as possible to minimize the effects of not having individual cell boundaries.

**Dot identification**

smFISH and SMOOTHY-FISH dots were detected within segmented cell regions using the previously described two-dimensional Gaussian wavelet-based detection and fitting procedure[23].

It is important to choose the appropriate thresholds for the dot detection, because it affects both FISH dot counts and colocalization. For example, in a two-channel image analysis, a too low threshold in the first channel picks up background noises as false FISH dots. The false FISH dots may colocalize to real FISH dots that don't have real colocalization in the second channel, making the colocalization fraction very high in the second channel. Since a lot of the false FISH dots in the first channel don't have colocalization, this makes the colocalization fraction very low in the first channel. On the other hand, a too high threshold in the first channel misses real FISH dots, causing the colocalization fraction very low in the second channel and very high in the first channel.

To avoid picking up background noise or missing real FISH dots, we set the threshold to the lowest while not picking up any single-pixel noise as FISH dots. For detection yield analysis, we set threshold = 30 for Alexa Fluor 488 SMOOTHY-FISH and threshold = 80 for ATTO647N smFISH (Fig. 3b). For detection yield analysis in tissues, we set threshold = 80 for Alexa Fluor 594 and threshold = 55 for ATTO647N smFISH (Fig. 3d). We confirmed that background noise was not picked up by negative control SMOOTHY-FISH probes (Supplementary Fig. 2). We set threshold = 30 for Alexa Fluor 488, threshold = 40 for Alexa Fluor 594, and threshold = 30 for ATTO647N in SMOOTHY-FISH probe brightness quantification (Supplementary Table 3). We set threshold = 30 for Alexa Fluor 488 and threshold = 40 for Alexa Fluor 594 in PKM isoform mapping (Fig. 4b). We set threshold =120 for Alexa 594 and threshold = 40 for ATTO647N in PKM isoform mapping in tissues (Fig. 4e). We set threshold = 30 for Alexa Fluor 488 in miRNA mapping (Fig. 5a, e). We set threshold = 80 for Alexa Fluor 594 in miRNA mapping in tissues (Fig. 5b - d). We set threshold = 100 for Alexa Fluor 488, threshold = 100 for Alexa Fluor 594, and threshold = 40 for ATTO647N in MS2-3-PP7-NiNTA conformation mapping (Fig. 6d); the high thresholds used were due to the stronger autofluorescence and dense SMOOTHY-FISH signals in the samples.

**Colocalization dot pairing**

Each candidate dot was analyzed within a 15 × 15 pixel window. For colocalization analysis, two dots from different fluorescence channels were considered colocalized if their peak-pixel centers were within 2.5 pixels[23] (1 pixel length = 108nm). When multiple dots were found within this distance threshold, the closest dot was assigned as the colocalized partner. Colocalization was performed as one-to-one pair matching, so each dot could be assigned to at most one colocalized dot in the other channel.

**Detection yield calculation**

Conventional smFISH was used as a practical benchmark for detection yield because each target transcript is robustly detected by the collective signals from a set of singly labeled probes (Fig. 1c). We evaluated this benchmark using two common smFISH probe-set designs for SDHA mRNA: ② two probe sets separately targeting upstream and downstream regions of the transcript and ③ two interleaved probe sets targeting alternating positions across the transcript. We measured ① SMOOTHY-FISH's detection yield at sites P07, P17, and P26 using the same approach (Fig. 3b).

We calculated the detection yield for individual cells as:

$$individual\ yield = \frac{2C}{A + B}$$

where C is the number of colocalized dots within the cell, A is the number of channel-1 FISH dots, and B is the number of channel-2 FISH dots. The mean ± SD of the individual yields from 50 cells are used to represent the detection yield of ① SMOOTHY-FISH and ②③ benchmarks.

This detection yield calculation also mitigated the bias from threshold determination in FISH dot detection. Inappropriate thresholds, either too low or too high, always cause extremely high colocalization in one channel and extremely low colocalization in another channel. Compared to other measurements (e.g. linear approximation: C = kA or C = kB, where slope k represents the yield) that calculate the yield based on one of the two channels, this measurement considered two channels and resulted in one number to represent the detection yield.

**Supplementary Note 2: SMOOTHY-FISH probe brightness quantification**

**Dot intensity fitting**

After dot positions were identified in each fluorescence channel, a window centered on each detected dot was selected from the original image for intensity fitting using the same MATLAB pipeline[23].

This pipeline fits fluorescent dots using a rotated asymmetric two-dimensional Gaussian model in two stages. In the first stage, candidate dots are iteratively fitted with two-dimensional Gaussian intensity distributions and subtracted from the image. This process is repeated until the intensity of the next candidate dot falls below a defined threshold, which was set to 10% of the integrated intensity of the first fitted dot in this analysis. This step generates an initial set of potentially overlapping Gaussian objects. In the second stage, the original image is refitted as a linear combination of the identified Gaussian objects, with their positions constrained near the values estimated in the first stage.

**Probe brightness quantification**

The brightness of each SMOOTHY-FISH probe design was represented by the median integrated intensity of all detected SMOOTHY-FISH dots for that design. We estimated the uncertainty in the median brightness by a bootstrap 95% confidence interval, because the number of detected dots varied substantially across all designs due to differences in fluorophore choice, fluorophore number, fluorophore spacing, and overall probe size (Fig. 2b).

For each probe design, bootstrap analysis was performed as follows. First, for a dataset containing N detected dots, N dots were randomly resampled with replacement, and the median integrated intensity of the bootstrap sample was calculated. This process was repeated 10,000 times to generate a bootstrap distribution of median intensities. The 95% confidence interval was then calculated from the 2.5th and 97.5th percentiles of the bootstrap distribution. Probe brightness quantification results are listed in Supplementary Table 3.

**Balance between brightness and yield**

As shown in Supplementary Table 3, 5-nt gap probes were brighter than 4-nt and 3-nt probes with the same number of fluorophores, and with the same gap size, brightness generally increased as the number of fluorophores increased, despite such a correlation was not strictly linear. However, as the number of FISH dots differ substantially across all probe designs, the behaviors of these probes were certainly affected by their gap sizes, the types of fluorophores, the number of fluorophores, and the total length.

So, we measured the yields of 5-nt gap probes with 3 to 10 Alexa Fluor 488 incorporated using the same approach described in Supplementary Note 1. The yield was the highest with 5 Alexa Fluor 488 in the 5-nt gap probe (Fig. 2c). And, the yield dropped drastically with 8 or more Alexa Fluor 488 in 5-nt gap probes, because non-specific probe retention occurred in these probe designs (Fig. 2c). Probe aggregation was seen in 4-nt gap probes with 8 or more Alexa Fluor 594 (Supplementary Table 8). Therefore, we decided to use 5-nt gap probes with 5 fluorophores (Alexa Fluor 488, Alexa Fluor 594, and ATTO647N) for all other experiments due to the balance between brightness and yield.

#### Supplementary Note 3: PKM1/2 isoform mapping

##### Target choice and mapping

We chose PKM1/2 as a target to demonstrate SMOOTHY-FISH's ability to map isoforms, because PKM1/2 has two mutually exclusive alternative spliced isoforms, which are well-investigated in previous studies[33,34]. PKM1 contains exon 9; PKM2 contains exon 10 (Figure 4a). The two exons have the same length of 167-nt and are 73% identical in sequence. Such a short sequence of differences highlighted the advantage of our single-probe mapping strategy.

To map the spliced transcripts, we designed SMOOTHY-FISH probes targeting the exon 8–9 and exon 8–10 junctions rather than within exon 9 and 10. The distinction between these two isoforms relied on the 2-nt difference at the 5' end of exon 9 and 10 (Fig. 4b). It was necessary to place flankers adjacent to the target sites to enhance the accessibility since RNA secondary structures were common near splice junctions. We performed PKM1/2 isoform mapping in cell lines twice with flipped fluorophore assignments on SMOOTHY-FISH probes to control for potential fluorophore-dependent bias in isoform quantification (Supplementary Fig. 4).

##### PKM analysis in HEK293 and SH-SY5Y cells

We measured the fraction of PKM2 within the total PKM expression in cell lines by linear regression (y = kx; where y = PKM2, x = total PKM, and slope k = fraction) (Fig. 4e).

##### PKM analysis in Mouse Brain Tissues

For analyzing that ratio of PKM1 to PKM2 in tissue samples, the number of each isoform was determined for each segmented whole tissue area. Instead of calculating the total ratio for each segmented area, the total number of each isoform was added across all segmented regions and used to determine the final percentage of each isoform.

**Mutually exclusive alternative splicing of PKM1/2 at tight ratio albeit high variation in total PKM expression in cell population**

SMOOTHY-FISH revealed that the splicing of PKM1/2 was maintained at a tight ratio despite substantial cell-to-cell variation in total PKM abundance within the cell population. This observation implies the PKM isoform ratio, rather than absolute transcript number, is critical for its cellular functions[52]. It is surprising that this ratio was so tightly regulated. How the cellular machineries contribute to such a tight PKM ratio across cells with different total PKM expression levels remains an interesting question. Alternative splicing is regulated through multiple cellular machineries, including spliceosome assembly[53,54], RNA-binding proteins[55,56], transcriptional kinetics[54], and chromatin context[54]. It is likely that a combination of multiple machineries cooperate to achieve this, as suggested by the previous studies[57].

**RT-PCR Analysis:**

Quantification of the PKM1 to PKM2 isoform ratio using RT-PCR and restriction enzyme digestion yielded two numbers: one ratio when digesting PKM1, and one ratio when digesting PKM2 (Supplementary Fig. 5). The percentage of PKM2 from this analysis is reported as the average of the two numbers by adding up the two percentages of PKM2, and dividing by 2. The resulting percentage was then converted into the final ratio of PKM1 to PKM2.

#### Supplementary Note 4: miRNA mapping in cell lines and mouse brain tissues

##### Target choice

We chose the miRNAs exhibiting different abundance in HeLa cells (miR-10a-5p, miR-16-5p, and miR-23a-3p)[58] to confirm SMOOTHY-FISH can map miRNAs in a large abundance range in cell line (Fig. 5a). To confirm this in tissues, we chose to map the hippocampus-enriched miR-218-5p in three different mouse brain tissue regions (Fig. 5b, c, d). We also confirm miR-218-5p localization in 3T3 cells (Fig. 5e).

##### miRNA analysis in HeLa and 3T3 cells

We measured the cytoplasmic fraction of these miRNAs in cell lines by linear regression ($y = kx$; where $y$ = cytoplasmic miRNAs of interest, $x$ = total miRNAs of interest, and slope $k$ = fraction) (Fig. 5f). We measured the nuclear and cytoplasmic concentration of these miRNAs in cell lines by linear regression ($y = kx + m$; where $y$ = nuclear or cytoplasmic miRNAs of interest, $x$ = nuclear or cytoplasmic area, $m$ = intercept, and slope $k$ = fraction) (Supplementary Fig. 9a, b). Though the intercept is extrapolated because cytoplasm area can not be zero, its presence in the linear fitting suggested a basal level of miRNA transcript number in the cytoplasm.

##### miRNA analysis in Mouse Brain Tissues

Whole area segmentation was done as previously described. In addition, nuclei were segmented carefully around the DAPI signals that were contained within the whole area segmentation. Dots were counted in both the whole area segmentation, as well as within the nuclei segmentation. Total dots were added for the nuclei segmentation to determine the total number of miRNA within all nuclei, and this value was subtracted from the total number of miRNA captured within the whole area segmentation to determine the number of miRNA found within the cytoplasmic area. The total number of dots in the nuclear area and cytoplasmic area was then used to determine the miRNA percentage of each subcellular region.

**miRNAs predominantly localized to cytoplasm at a tight cytoplasm-to-nucleus ratio**

SMOOTHY-FISH mapping revealed that the mapped miRNAs predominantly localized to the cytoplasm, with a tightly consistent cytoplasmic fraction across cells exhibiting different total miRNA abundances. miRNAs mature in cytoplasm[59] and perform regulatory functions in cytoplasmic mRNA destabilization and translational repression[60,61]. Our observation implies that cells may actively regulate a tight cytoplasm-to-nucleus ratio for the functions of miRNAs. Previous work by Wei et al. (2014) demonstrated that Importin-8 facilitates the nuclear import of mature miRNAs through interactions with Argonaute 2 (Ago2)[62]. Whether this nuclear import pathway or other mechanisms contribute to maintaining relatively consistent nuclear–cytoplasmic miRNA distributions across cells with different total miRNA abundances remains to be investigated.

**miRNAs maintained a relatively stable concentration in cytoplasm**

SMOOTHY-FISH quantification revealed positively linear correlations between cytoplasmic miRNA abundance and cytoplasmic area across individual cells, suggesting that cells maintain relatively stable miRNA concentration despite difference in cytoplasm area. Interestingly, a similar scaling relationship was previously reported by Padovan-Merhar et al. (2015), who demonstrated that mRNA abundance increases with cell volume through changes in global transcriptional output, contributing to the maintenance of relatively stable transcript concentrations across cells of different sizes[63]. Because miRNAs regulate cytoplasmic mRNAs, the relative abundance of miRNAs and their available mRNA target sites can influence the extent of miRNA-mediated repression[64]. This observation suggests that scaling of miRNA and mRNA abundance with cell size may help maintain the balance between miRNAs and their mRNA targets, achieving consistent mRNA repression.

#### Supplementary Note 5: MS2-3-PP7-NiNTA conformational mapping

##### Target choice

We chose the MS2-3-PP7-NiNTA structural element for mapping because it was well-studied in previous work[65]. It consists of a basal stem and three hairpins (Fig. 6a), of which the combinations provide multiple conformation for SMOOTHY-FISH to map its in situ conformation state.

##### Defining the seven conformations for mapping

The minimum free energy structure and centroid structure predicted by Vienna RNAfold[30] both corresponded to Conformation *I* (Fig. 6a). Additional conformations were defined by combinations of folded and unfolded states of the basal stem, MS2 hairpin, PP7 hairpin, and NiNTA hairpin that generated accessible and distinct junction sequences (Supplementary Fig. 11). Each additional conformation was mapped together with Conformation *I* to test whether it could coexist with the predicted most stable state and to reduce potential bias from probe-induced stabilization of a single conformational state (Supplementary Fig. 12).

##### Flanker choice

Flanking oligonucleotides were designed to bind immediately adjacent to the common-region site to maximize its accessibility, because common-region signal was used to estimate the total transcript population. For all conformational mapping experiments, flanking oligonucleotides were also designed to bind immediately adjacent to the Conformation *I* junction sequence. This design was used to mitigate the neighboring RNA sequences' influences on MS2-3-PP7-NiNTA folding and junction accessibility.

##### Calculation of conformational fractions

For each conformational state, the estimated fraction was calculated as the fraction of common-region signals that colocalized with the corresponding conformation-specific junction signal. When Conformation *I* and an additional conformation were mapped together, each conformational fraction was calculated independently using the common-region signal as the reference for the total transcript population. Colocalization fractions are presented as median with 12.5th and 87.5th percentile range.

**Junction specificity**

Some junction sequences had imbalanced GC content between the two sides of the junction, raising the possibility that a junction probe could bind only one side of the target and generate false-positive signals. To test whether junction detection required recognition of both sides of the junction, half-junction control probes were designed to target either the upstream or downstream 10-nt sequence of the junction (Supplementary Fig. 10). This control was performed for Conformations *I* and *V*, because these two conformations coexist at high fractions. The lack of detectable signal from half-junction probes supported the specificity of full junction-based detection.

**Conformational heterogeneity in situ and comparison with RNA structure prediction**

SMOOTHY-FISH revealed that MS2-3-PP7-NiNTA populated two major conformational states in cells: Conformation *I*, predicted by RNAfold as both the minimum-free-energy and centroid structure, and Conformation *V*, in which only the MS2 hairpin is folded (Fig. 6d,e). The coexistence of these two structurally distinct states at comparable estimated fractions reveals substantial conformational heterogeneity beyond the single most stable structure predicted in silico. Similar structural heterogeneity has been reported in previous studies, including the identification of alternative RNA conformations within HIV-1 transcripts[66], supporting that RNA molecules can populate ensembles of distinct conformational states[67].

Interestingly, Conformations *V* and *VI* both contain only one folded hairpin, MS2 and PP7, respectively, yet their estimated fractions differed substantially. While Conformation *V* represented approximately 18% of the transcript population, Conformation *VI* accounted for less than 1%. For the Conformation *V* junction to become accessible, the basal stem and PP7 hairpin must unfold, whereas accessibility of the Conformation *VI* junction requires unfolding of the basal stem, MS2 hairpin, and NiNTA hairpin. These different structural requirements raise the possibility that the folding of individual hairpins is influenced by interactions with neighboring hairpins and the basal stem, rather than solely by their intrinsic stability.

Apart from local structural interactions, previous work has also demonstrated that RNA folding in cells can be influenced by ATP-dependent remodeling and other cellular factors[68]. How these cellular factors and local structural interactions contribute to the observed conformational distribution remains an open question.

In total, the conformational states of an RNA structure can behave differently in cells than in silico modeling. Many RNA structural conformations and their functions remain to be discovered within their intact cellular environment. SMOOTHY-FISH offers a new way to reveal in situ RNA structure ensembles and provides opportunities to investigate the related cellular factors through spatial mapping. The newly discovered in situ RNA conformations can provide further feedback to the in silico modeling for better and more extensive predictions.

**Limitations**

When Conformation *I* was mapped together with Conformation *V*, the estimated Conformation *I* fraction was slightly lower than when Conformation *I* was mapped with other alternative conformations (Supplementary Fig. 12). One possible explanation is that the Conformation *V* probe stabilized a small subset of transcripts that converted slowly between Conformation *I* and *V* states, thereby shifting the apparent conformational distribution. This potential probe-induced effect does not affect the conclusion that Conformations *I* and *V* coexist in cells, because the combined fraction of Conformations *I* and *V* was substantially higher than the combined fraction when Conformation *I* mapped in experiments paired with other alternative conformations.

#### Supplementary Note 6: SDHA mRNA mapping using the initiator design

##### SMOOTHY-FISH using the initiator design

In the initiator-based design, a target-specific initiator oligonucleotide was used to bridge the RNA target site and a universal readout probe (Supplementary Fig. 13a). An initiator contained a 20-nt target-binding region, a 20-nt readout-probe-binding sequence, and a flexible linker. The readout-probe-binding sequence was designed to be orthogonal to the human transcriptome to minimize off-target hybridization. To assess detection yield at the SDHA mRNA P26 site, we used an initiator targeting the P26 site together with the same SDHA smFISH probes (Supplementary Fig. 13c). With or without flanking oligonucleotides, the initiator-based design achieved detection yields approaching the practical benchmark established by conventional smFISH (Supplementary Fig. 13d). In contrast, the universal readout probe produced minimal signal in the absence of the initiator (Supplementary Fig. 13b), supporting initiator-dependent target detection.

##### Modified SMOOTHY-FISH workflow

For the initiator-based design, the initiator hybridization, post-initiator-hybridization washes, and SMOOTHY-FISH universal readout probe hybridization steps were added between the post-digestion washes and the post-hybridization washes step. After post-digestion washes, samples were hybridized with 20 nM initiator oligonucleotide and 0.5 ng/μL total smFISH probes in hybridization buffer supplemented with 1% v/v murine RNase inhibitor for ≥16 h at 30°C. After initiator hybridization, samples were washed with wash buffer (2× SSC, 20% formamide, 0.1% Tween-20) at 30°C for 15 min and washed with 2× SSC for 15 min at room temperature on a benchtop shaker. Samples were equilibrated in wash buffer for 5 min. Hybridization buffer containing 1% v/v murine RNase inhibitor, 20 nM each SMOOTHY-FISH probe was added to each well. Samples were incubated at 30°C for 4 h. Samples were then processed through the standard SMOOTHY-FISH post-hybridization and imaging workflow.

#### Supplementary Note 7: Technical considerations and potential extension of SMOOTHY-FISH

##### Single-probe specificity and detection considerations

Because SMOOTHY-FISH relies on hybridization through an approximately 20-nt target-binding region, we implemented multiple measures to optimize detection specificity. Hydrogel embedding and sample clearing were used to remove cellular proteins and lipids that could contribute to nonspecific probe binding and to reduce molecular crowding and nonspecific probe retention. We also used BLAST to screen candidate target sequences against the transcriptome to minimize potential off-target hybridization.

Nevertheless, off-target hybridization remains possible despite these measures. BLAST-based screening primarily identifies sequence similarities within annotated linear transcript sequences and does not fully account for RNA structures formed in situ. For example, RNA folding may bring nonadjacent sequences into proximity, potentially generating unintended junction motifs that resemble the intended structural target. Incorporating experimentally derived RNA structural information into target-site screening could provide an additional means of reducing such off-target interactions.

We introduced flanking oligonucleotides to improve single-probe hybridization efficiency, as local target accessibility varies along an RNA transcript and can influence detection yield. Although limited accessibility presents a challenge for efficient detection, differences in probe hybridization may also provide information about local RNA environments[69]. For example, the P07 site of SDHA mRNA exhibited a lower detection yield without flanking oligonucleotides than the P17 and P26 sites, potentially reflecting differences in local target accessibility. Such measurements could provide complementary information for investigating RNA folding and RNA–protein interactions.

Another consideration arises when the same short target motif is present in multiple transcript species. Because a single SMOOTHY-FISH probe recognizes only its target sequence, it cannot distinguish transcripts that contain identical target motifs but differ in surrounding sequences. One potential strategy is to introduce a second SMOOTHY-FISH probe targeting a neighboring transcript-specific region, enabling transcript identity to be assessed through signal colocalization. Alternatively, sequence-specific RNA anchors could potentially enrich the retention of selected transcripts within the hydrogel, reducing interference from other RNA species carrying the same motif.

Finally, conventional smFISH probe sets were used as a reference for evaluating SMOOTHY-FISH detection yield and establishing a practical benchmark. However, given the improvements in target accessibility achieved using flanking oligonucleotides, hybridization of multiple smFISH probes along the transcript might also alter local RNA accessibility[70] and influence the measured detection yield of SMOOTHY-FISH. If such an effect exists, hybridization of additional oligonucleotides to regions surrounding the target motif could potentially improve single-probe detection efficiency.

**Hydrogel preparation and RNA retention**

A key consideration in hydrogel preparation was maintaining a sufficiently thin gel layer (≤1 mm) to facilitate probe diffusion and hybridization. In preliminary experiments using thicker hydrogels (up to 5 mm), both SMOOTHY-FISH and conventional smFISH probes failed to produce substantial signals, suggesting limited probe penetration and hybridization. Therefore, thin hydrogels were used throughout the SMOOTHY-FISH workflow to maintain efficient target detection.

Although poly(dT) anchors were used to retain mRNAs within the hydrogel, we observed substantial retention of long RNAs even without anchoring, potentially because their large molecular sizes limited diffusion out of the hydrogel matrix. If removal of unanchored RNAs is desired, alternative hydrogel formulations with larger effective pore sizes could potentially facilitate RNA diffusion. For example, modifying the acrylamide concentration or the ratio of bis-acrylamide to acrylamide could alter the gel mesh size. Expansion-based hydrogel formulations may also provide an alternative approach.

In contrast, small RNAs required anchoring for efficient retention within the hydrogel, as demonstrated in Fig. 5. During SMOOTHY-FISH development, we evaluated two small RNA anchoring strategies: (1) poly(A) tailing followed by poly(dT) anchors hybridization and (2) poly(5-vinyl-U) tailing using modified 5-vinyl-UTP. For the second strategy, we attempted to extend the 3′ ends of small RNAs with 5-vinyl-uridine residues through template-independent nucleotide addition by the terminal uridylyltransferase Cid1. The introduced vinyl groups were intended to enable direct covalent incorporation of the modified RNA tails into the polyacrylamide matrix during gel polymerization. Although SMOOTHY-FISH detected some signals from 24-nt target RNA oligonucleotides processed using this approach, PAGE analysis indicated that only a small fraction of the RNA substrates were successfully extended. This low tailing efficiency may reflect limited compatibility between Cid1 and the modified 5-vinyl-UTP. Given the inefficient RNA extension and limited detection performance, we selected poly(A) tailing followed by poly(dT)-mediated anchoring as the small RNA retention strategy for the optimized SMOOTHY-FISH workflow.

**Considerations for tissue applications**

Modifications to the SMOOTHY-FISH workflow for tissues were mainly done for initial sample handling, permeabilization, and digestion. Mouse brain tissue samples needed to be fixed prior to cryopreservation in order to be stored as free-floating sections in 70% ethanol. Utilizing free-floating tissue samples allows for easier sample handling and minimizing consumable usage, but may not be possible for other tissue types.

Initial testing was done for alternative permeabilization reagents, such as SDS to improve anchor and probe accessibility, but this usually led to a noticeable loss of RNA (Data not shown). SDS permeabilization also resulted in the free-floating tissue samples to lose more structure, and were subsequently more difficult to work with prior to gelation (Data not shown).

A double digestion was performed for the mouse brain tissue samples due to samples appearing not fully cleared before probe hybridization, contributing to higher background signal (Data not shown). We theorized that both the protein digestion and the lipid removal were not fully complete, and decided to optimize them in two separate steps. Higher SDS concentrations help promote lipid removal but begin to inhibit proteinase K, while proteinase K runs more optimally at 0.5% SDS. Washes in-between digestions were done to remove any digested lipids and proteins from that round.

**Potential extensions of SMOOTHY-FISH**

SMOOTHY-FISH could be adapted to incorporate additional fluorophores beyond the three tested in this study, potentially increasing its multiplexing capacity. This would be particularly beneficial for tissue applications, where strong green autofluorescence in mouse brain tissue limited our imaging to Alexa Fluor 594 and ATTO647N. Expanding the range of compatible fluorophores, particularly in the red and far-red spectral regions, could enable additional imaging channels and improve multiplexed RNA detection in tissue samples.

Another potential extension is to incorporate expandable hydrogels into the SMOOTHY-FISH workflow in place of the current non-expandable hydrogel. Expansion microscopy enables higher effective spatial resolution by physically separating neighboring RNA molecules[71,72]. Expansion-based sample processing might also facilitate the removal of unanchored long RNAs, potentially enabling more selective RNA retention when combined with transcript-specific anchoring strategies. However, several technical considerations would need to be addressed, particularly hydrogel immobilization and probe diffusion. In the current workflow, hydrogels are immobilized on silanized glass surfaces through covalent attachment, which would restrict physical expansion if maintained during swelling. An expandable hydrogel would therefore need to be released from the glass surface before expansion and could subsequently be stabilized through re-embedding in a non-expandable hydrogel. The resulting gel thickness and composition would also need to be optimized to maintain efficient SMOOTHY-FISH probe penetration and hybridization in expanded samples.

### Supplementary Figures:

**Supplementary Figure 1. Examples of non-optimal probe designs.**

**a**, Probe designs with too small spacing between each fluorophore failed to generate detectable fluorescent dot signals even with many fluorophores. **b**, Probe designs with too few fluorophores generated vague signals that were difficult to be distinguished from the background. **c**, Probe designs with too large in size caused non-specific probe retention in the cells.

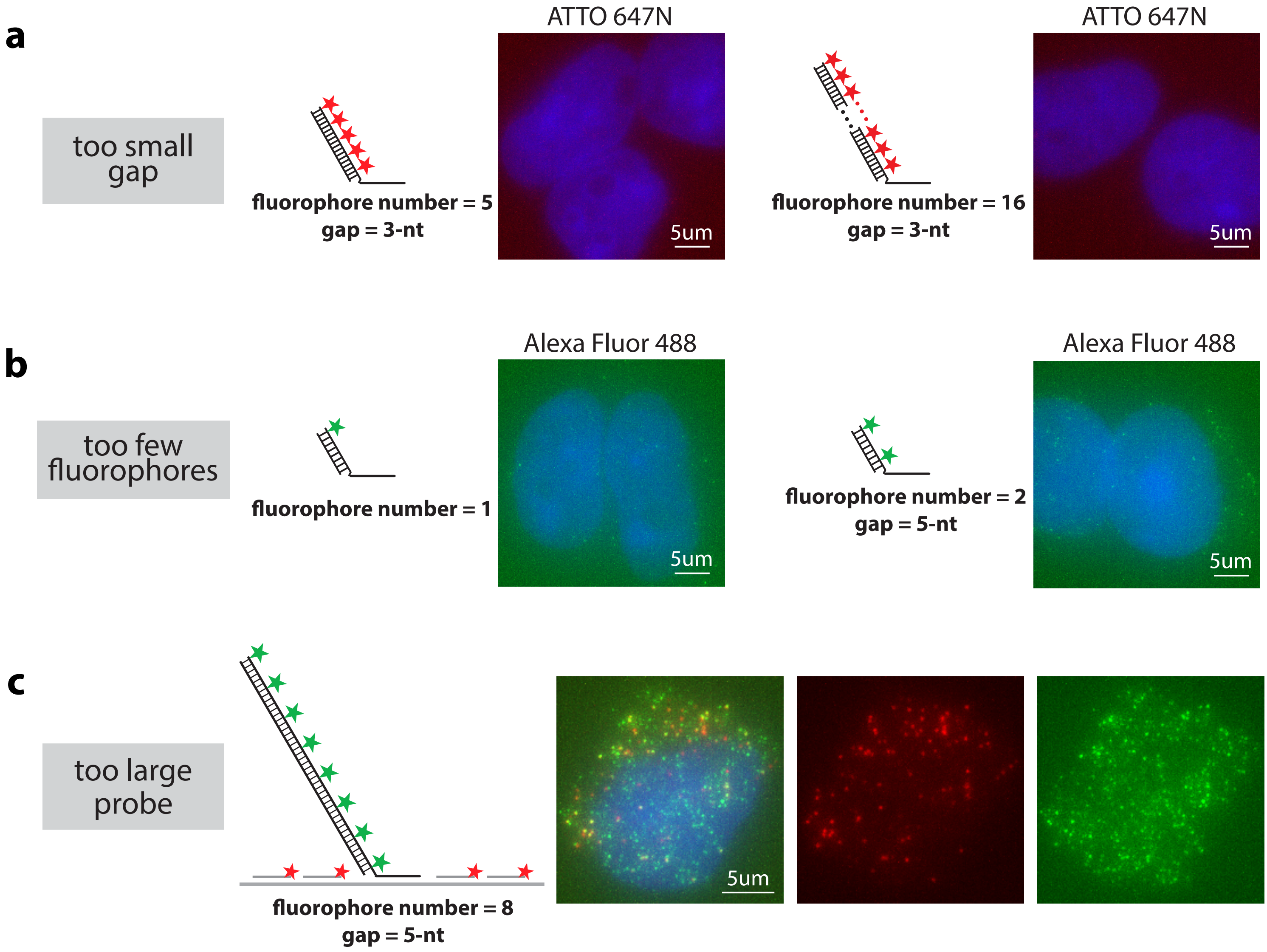
a
too small gap
fluorophore number = 5
gap = 3-nt
ATTO 647N
5um
fluorophore number = 16
gap = 3-nt
ATTO 647N
5um
b
too few fluorophores
fluorophore number = 1
Alexa Fluor 488
5um
fluorophore number = 2
gap = 5-nt
Alexa Fluor 488
5um
c
too large probe
fluorophore number = 8
gap = 5-nt
5um

**Supplementary Figure 2. SMOOTHY-FISH specificity validation by negative control probes.**

Negative control mapping by **a**, a NULL SMOOTHY-FISH probe (green) lacking the single-stranded targeting region; **b**, an orthogonal SMOOTHY-FISH probe (green) targeting a sequence orthogonal to human transcriptome in HEK293 cells. SDHA smFISH probes (red) were used to confirm the absence of SMOOTHY-FISH signal was not due to RNA degradation. SMOOTHY-FISH and smFISH dot count per cell were represented by mean ± SD (n = 50 cells) in the bar graph.

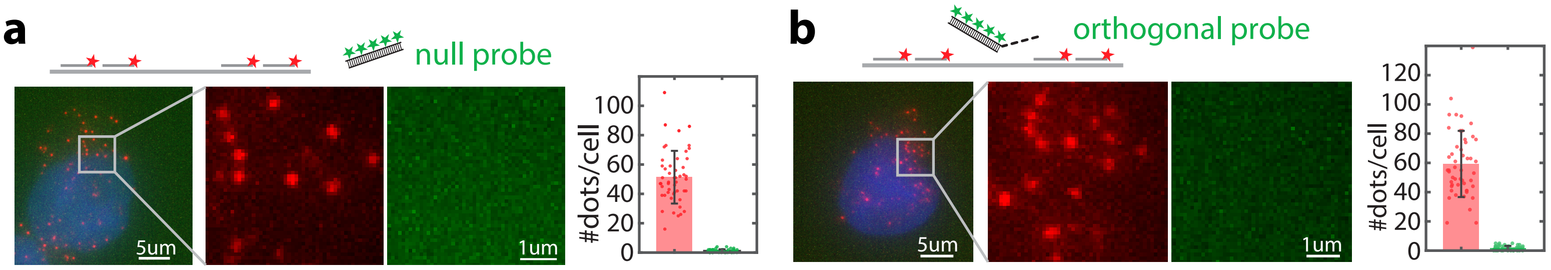
a
null probe
5um
1um
#dots/cell
0
20
40
60
80
100
b
orthogonal probe
5um
1um
#dots/cell
0
20
40
60
80
100
120

**Supplementary Figure 3. SMOOTHY mapped PKM1 and PKM2 isoform expression ratio in SH-SY5Y cells.**

SMOOTHY-FISH distinguished PKM1 (green) and PKM2 (red) in SH-SY5Y cells. Zoomed image insets showed regions with different total PKM expression.

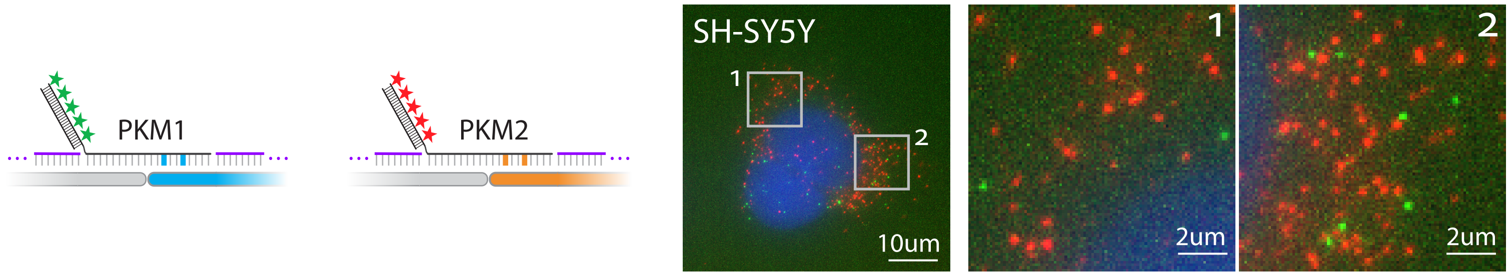
PKM1
PKM2
SH-SY5Y
1
2
10um
1
2um
2
2um

**Supplementary Figure 4. SMOOTHY-FISH mapped PKM1 and PKM2 isoform expression ratios in cell lines and mouse brain tissue with flipped fluorophore usage.**

PKM isoform mapping was not biased by fluorophore usage in SMOOTHY-FISH probes. The same PKM1 (red) and PKM2 (green) isoform mapping experiment as in **Fig. 4** and **Supplementary Fig. 3** was repeated with flipped fluorophore usage. Zoomed image inset showed cells with different total PKM expression.

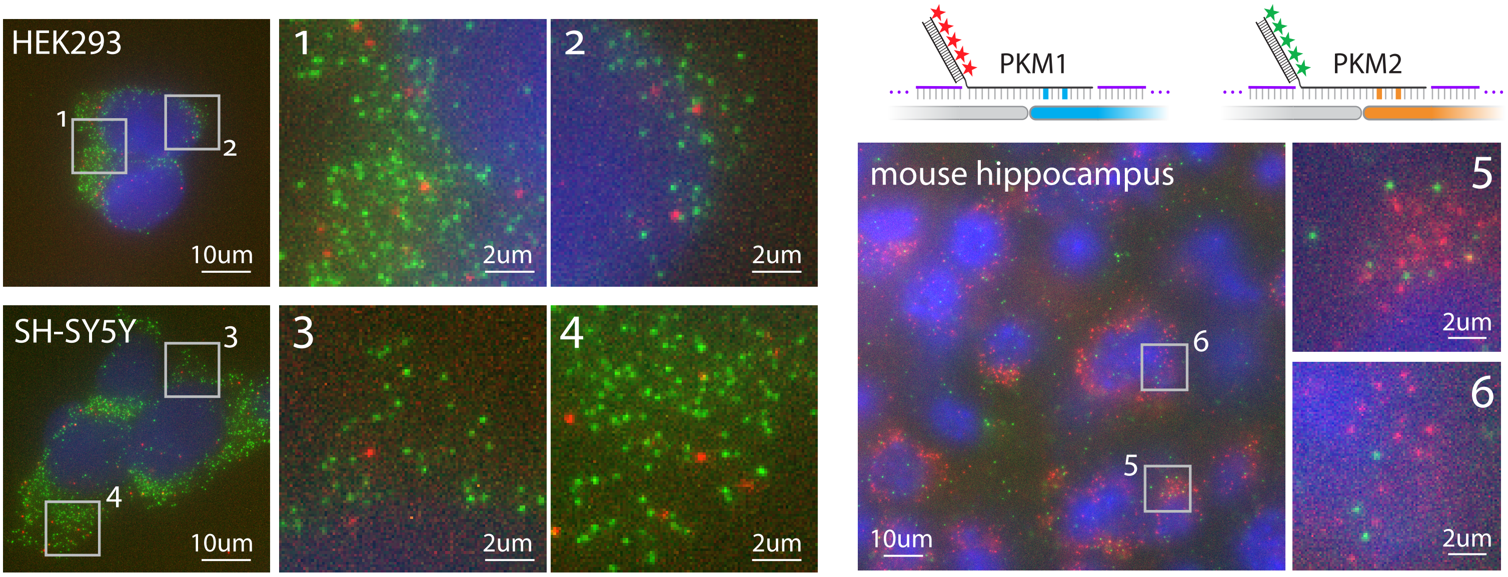

HEK293
1
2
10um
2um
2um
SH-SY5Y
3
4
10um
3
2um
4
2um
PKM1
PKM2
mouse hippocampus
6
5
10um
5
2um
6
2um

**Supplementary Figure 5. PKM1 and PKM2 isoform expression ratios in cell lines and mouse brain tissue were confirmed by RT-PCR with restriction digestion.**

PKM1 and PKM2 isoform expression ratios in **a**, HEK293 cells, **b**, SH-SY5Y cells, and **c**, mouse brain tissue were confirmed by running electrophoresis of the restriction digested PKM-specific RT-PCR amplicons in 1% agarose gel. RT-PCR amplicon sizes were indicated in the gel images. PsiI-v2 specifically digested human PKM1 and mouse Pkm1 amplicon; PstI-HF specifically digested PKM2 amplicon; SfoI specifically digested mouse Pkm2 amplicon. Percentages of PKM2 were indicated at the bottom of the gel image. RT-PCR primer sequences are listed in Supplementary Table 7.

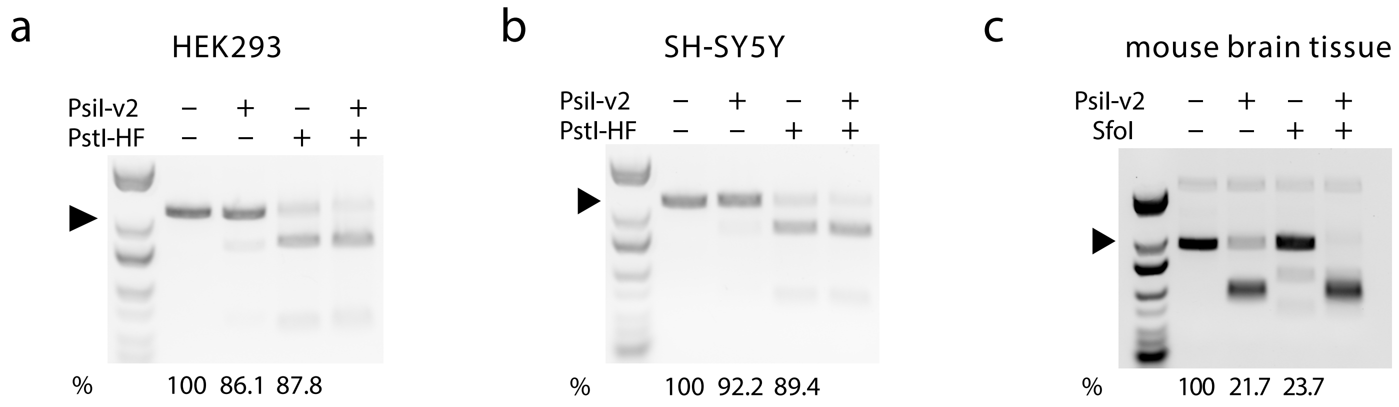

a
HEK293
Psil-v2 − + − +
PstI-HF − − + +
% 100 86.1 87.8
b
SH-SY5Y
Psil-v2 − + − +
PstI-HF − − + +
% 100 92.2 89.4
c
mouse brain tissue
Psil-v2 − + − +
SfoI − − + +
% 100 21.7 23.7

**Supplementary Figure 6. Similarly tight PKM1/2 isoform ratio by SMOOTHYY-FISH mapping with flipped fluorophore usage.**

PKM isoform mapping was not biased by fluorophore usage in SMOOTHY-FISH probes. The same PKM1 (red) and PKM2 (green) isoform mapping experiment as in **Fig. 4** and **Supplementary Fig. 3** was repeated with flipped fluorophore usage. The same linear fitting was performed as in **Fig. 4f**.

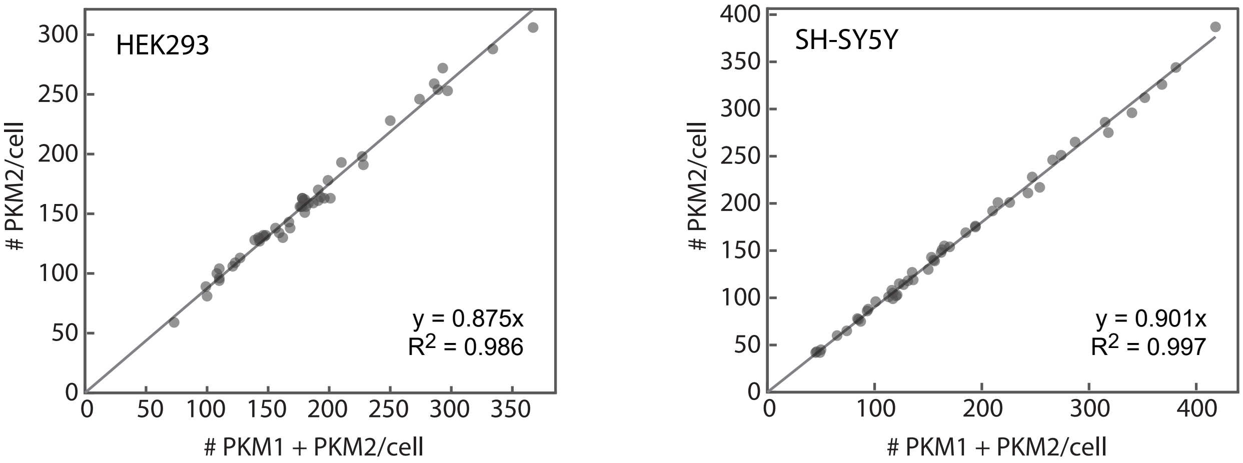
HEK293
y = 0.875x
R2 = 0.986
### PKM2/cell
### PKM1 + PKM2/cell
0
50
100
150
200
250
300
350
SH-SY5Y
y = 0.901x
R2 = 0.997
400

**Supplementary Figure 7. SMOOTHY-FISH quantitatively mapped miRNA abundance in cytoplasm, nucleus, and whole HeLa cells.**

**a**, SMOOTHY-FISH mapped miR-10a-5p, miR-16-5p, and miR-23a-3p in HeLa cells with and without the poly(A) tailing procedure. The abundance of these three miRNAs in **b**, the whole cell, **c**, the cytoplasm, and **d**, the nucleus of HeLa cells are shown as mean ± SD (n = 50 cells) in the bar graph. (Supplementary Note 4)

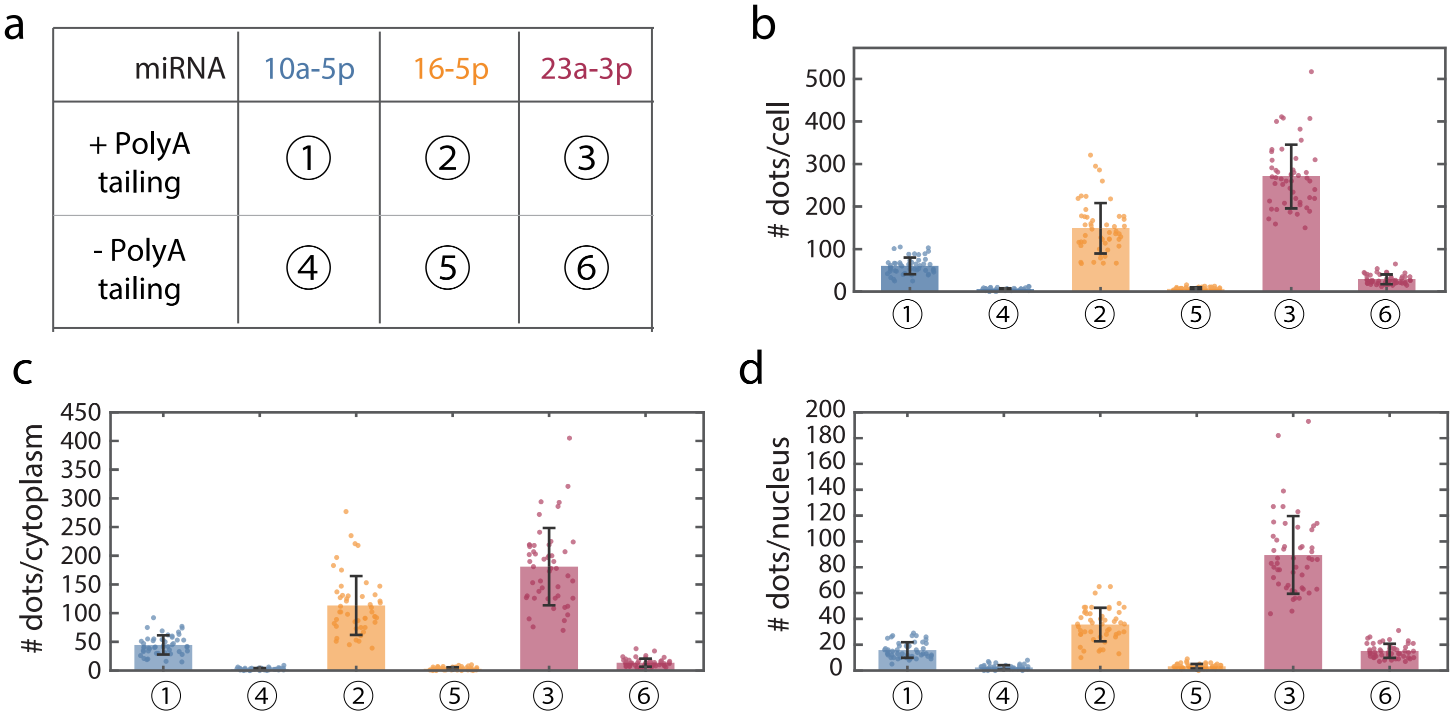

a
miRNA
10a-5p
16-5p
23a-3p
+ PolyA tailing
① ② ③
- PolyA tailing
④ ⑤ ⑥
b
### dots/cell
c
### dots/cytoplasm
d
### dots/nucleus
① ④ ② ⑤ ③ ⑥

**Supplementary Figure 8. SMOOTHY-FISH mapped miR-218-5p in three regions of the mouse brain tissue.**

SMOOTHY-FISH mapped miR-218-5p in mouse hippocampus, midbrain, and hypothalamus.

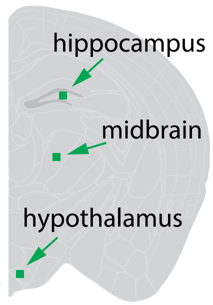

Map miRNA-218-5p
across mouse brain

**Supplementary Figure 9. SMOOTHY-FISH showed cytoplasmic miRNA abundance was strongly correlated to the cytoplasm area.**

Linear fitting of **a**, nuclear SMOOTHY-FISH dots vs. nucleus area and **b**, cytoplasmic SMOOTHY-FISH dots vs. cytoplasm area was calculated for miR-10a-5p, miR-16-5p, miR-23a-3p, and miR-218-5p (n = 50 cells). The linear correlations ($R^2$) in cytoplasm are stronger than in nucleus for all four mapped miRNAs.

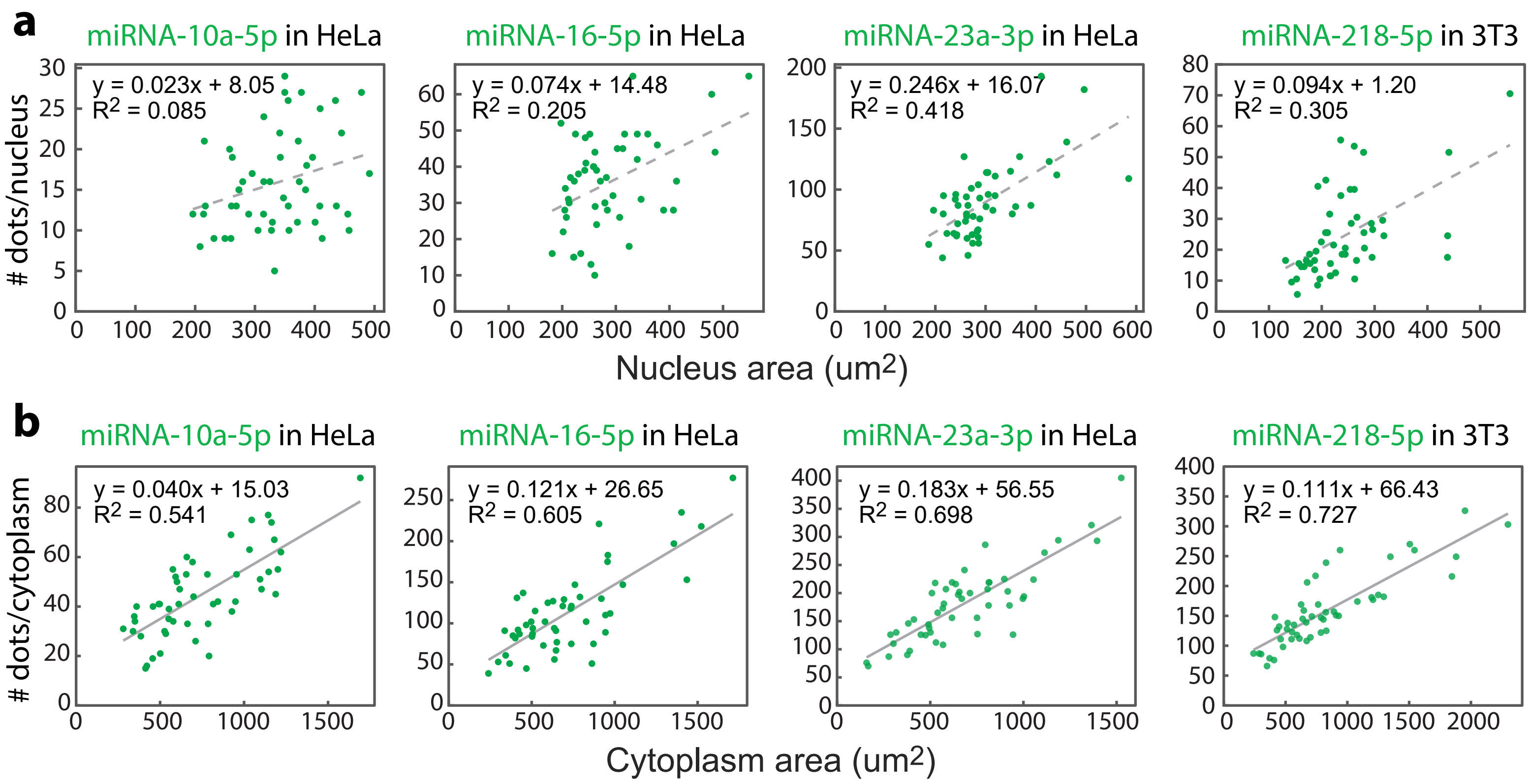
a
miRNA-10a-5p in HeLa
y = 0.023x + 8.05
R2 = 0.085
miRNA-16-5p in HeLa
y = 0.074x + 14.48
R2 = 0.205
miRNA-23a-3p in HeLa
y = 0.246x + 16.07
R2 = 0.418
miRNA-218-5p in 3T3
y = 0.094x + 1.20
R2 = 0.305
### dots/nucleus
Nucleus area (um2)
b
miRNA-10a-5p in HeLa
y = 0.040x + 15.03
R2 = 0.541
miRNA-16-5p in HeLa
y = 0.121x + 26.65
R2 = 0.605
miRNA-23a-3p in HeLa
y = 0.183x + 56.55
R2 = 0.698
miRNA-218-5p in 3T3
y = 0.111x + 66.43
R2 = 0.727
### dots/cytoplasm
Cytoplasm area (um2)

**Supplementary Figure 10. Half-junction SMOOTHY-FISH probes confirmed the specificity of MS2-3-PP7-NiNTA conformation mapping.**

Half junction SMOOTHY-FISH probe (red) targeted only the 10-nt sequence of **a**, conformation *I* upstream junction; **b**, conformation *I* downstream junction; **c**, conformation *V* upstream junction; and **d**, conformation *V* downstream junction. Sequences of the half-junction probes are listed in Supplementary Table 9.

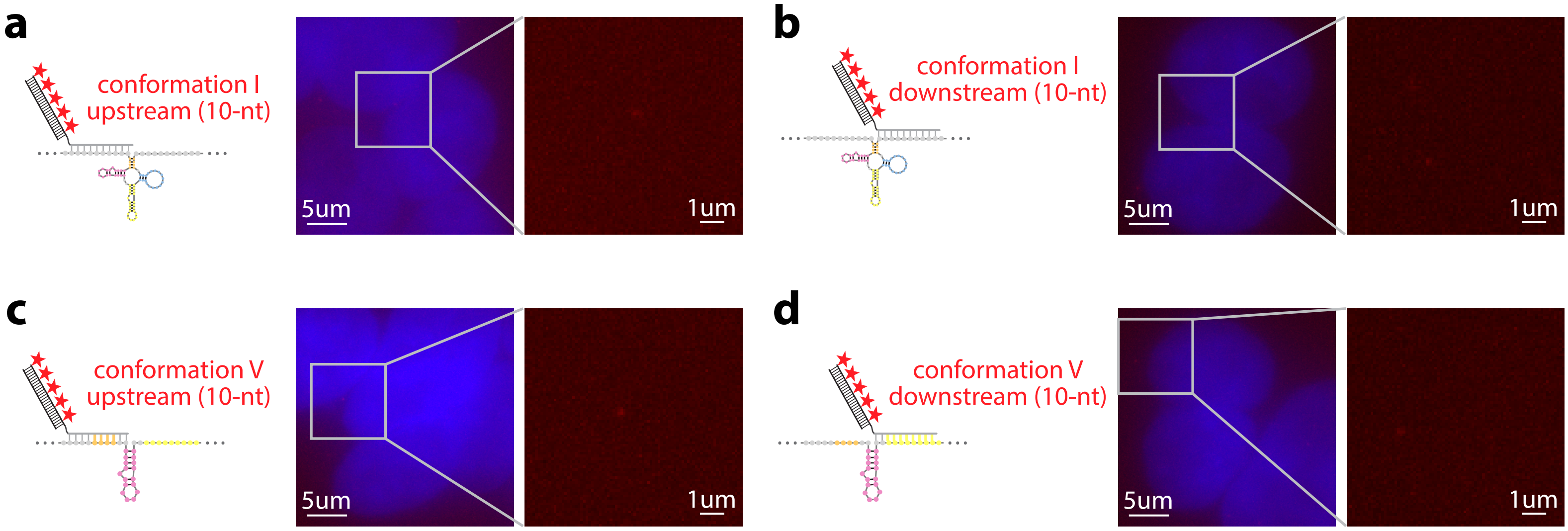

a
conformation I
upstream (10-nt)
5um
1um
b
conformation I
downstream (10-nt)
5um
1um
c
conformation V
upstream (10-nt)
5um
1um
d
conformation V
downstream (10-nt)
5um
1um

**Supplementary Figure 11. The RNA junctions created by seven possible MS2-3-PP7-NiNTA conformations.**

**a**, conformation *I*. **b**, conformation *II*. **c**, conformation *III*. **d**, conformation *IV*. **e**, conformation *V*. **f**, conformation *VI*. **g**, conformation *VII*. **h**, RNA junction target sequences created by each conformation.

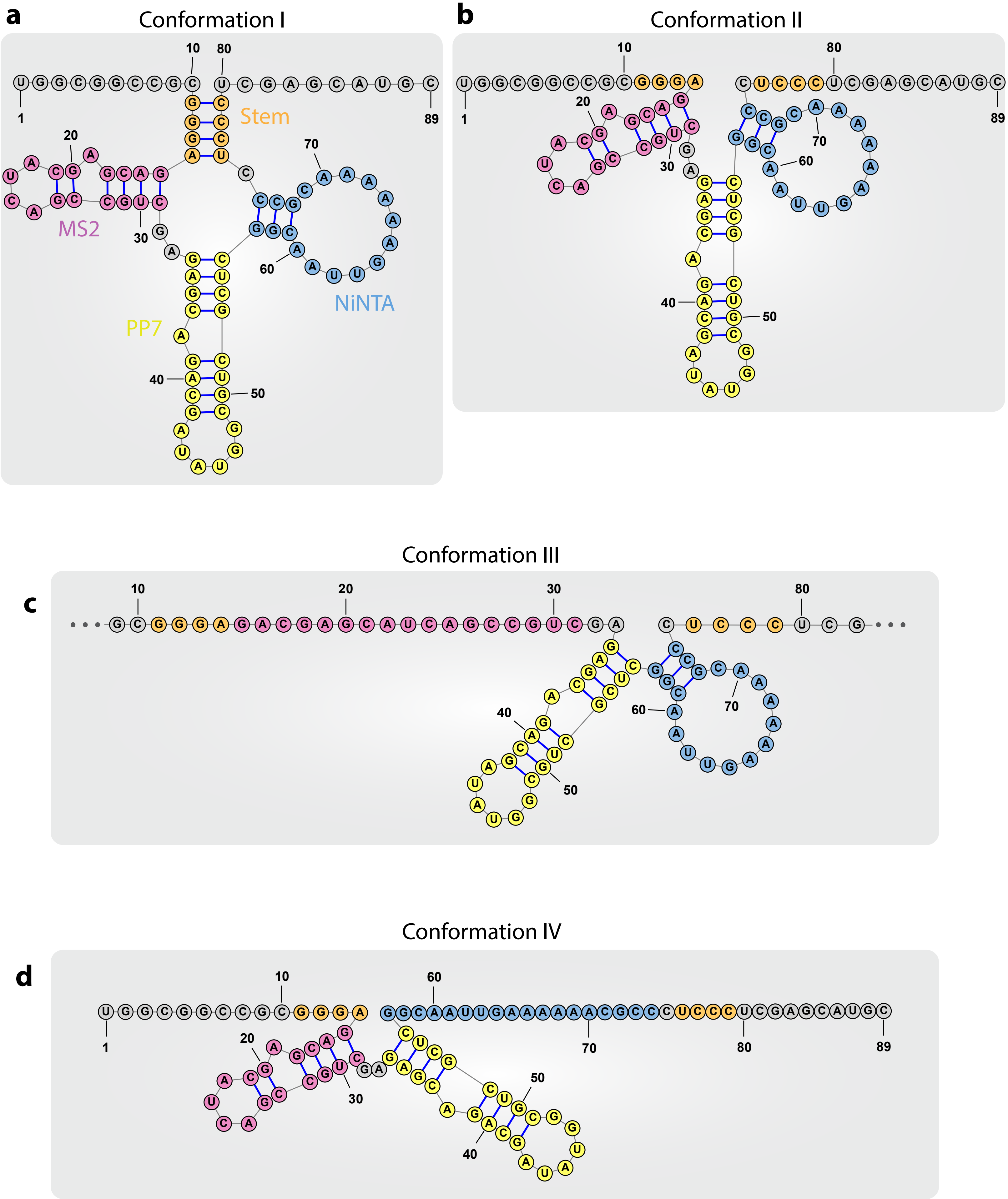
a
Conformation I
Stem
MS2
NiNTA
PP7
b
Conformation II
c
Conformation III
d
Conformation IV

**e**

Conformation V

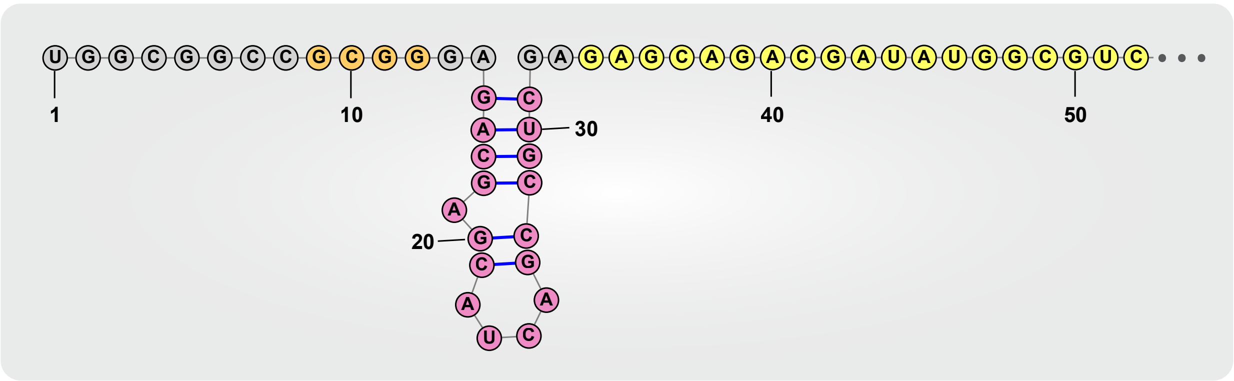

**f**

Conformation VI

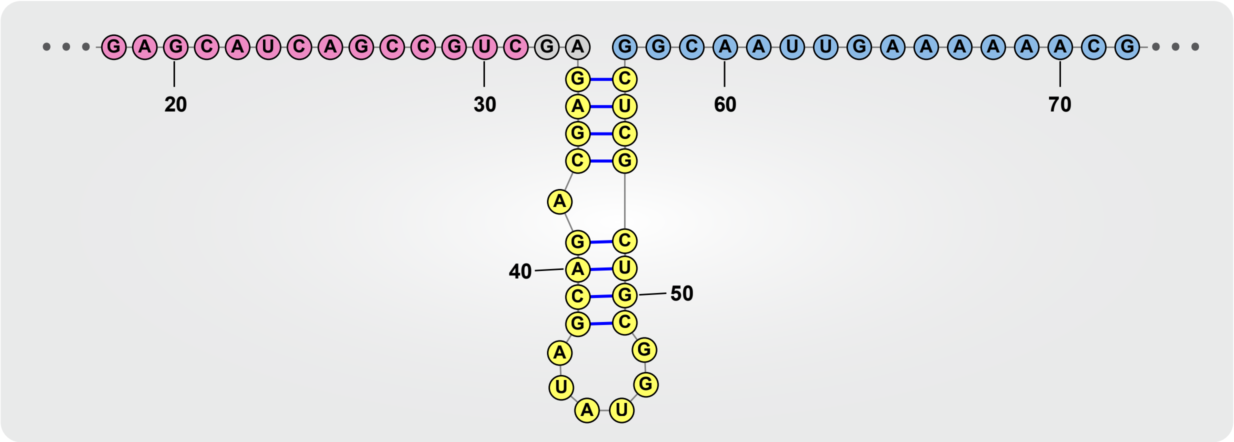

**g**

Conformation VII

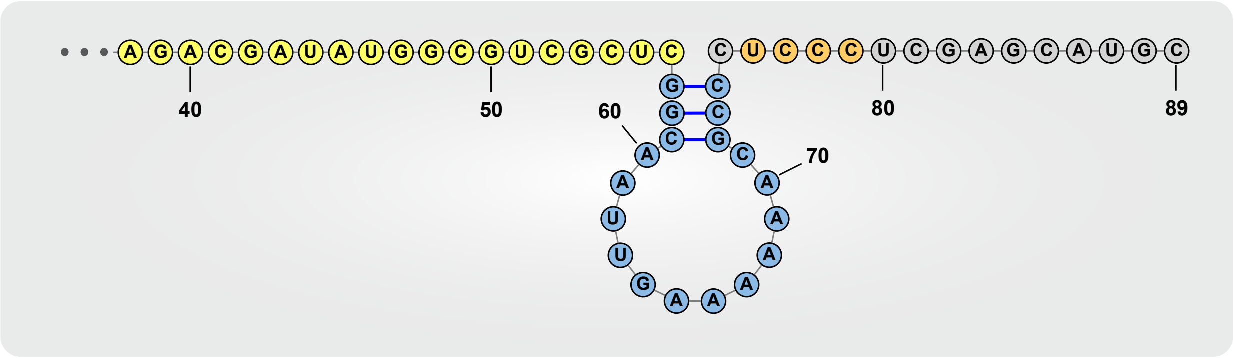

**h**

| | RNA junction target sequence (5'->3') |
|---|---|
| Conformation I | UGGCGGCCGCUCGAGCAUGC |
| Conformation II | GGCCGCGGACUCCCUCGAG |
| Conformation III | CAGCCGUCGACUCCCUCGAG |
| Conformation IV | GGCCGCGGAGGCAAUUGAA |
| Conformation V | GGCCGCGGAGAGAGCAGAC |
| Conformation VI | CAGCCGUCGAGGCAAUUGAA |
| Conformation VII | GGCGUCGCUCCUCCCUCGAG |

**Supplementary Figure 12. SMOOTHY-FISH mapped the co-existent fraction of MS2-3-PP7-NiNTA conformation I and each alternative conformation.**

Three-colored SMOOTHY-FISH mapping of the common region (green), conformation *I* junction (red), and **a**, conformation *II* junction (cyan); **b**, conformation *III* junction (cyan); **c**, conformation *IV* junction (cyan); **d**, conformation *VI* junction (cyan); and **e**, conformation *VII* junction (cyan). Colocalized dots between the common region and conformation I junction are circled in yellow. The fraction of each conformation is shown as median with the 12.5th–87.5th percentile ranges (n = 50 cells) in the bar graph. See the calculation in Supplementary Note 5.

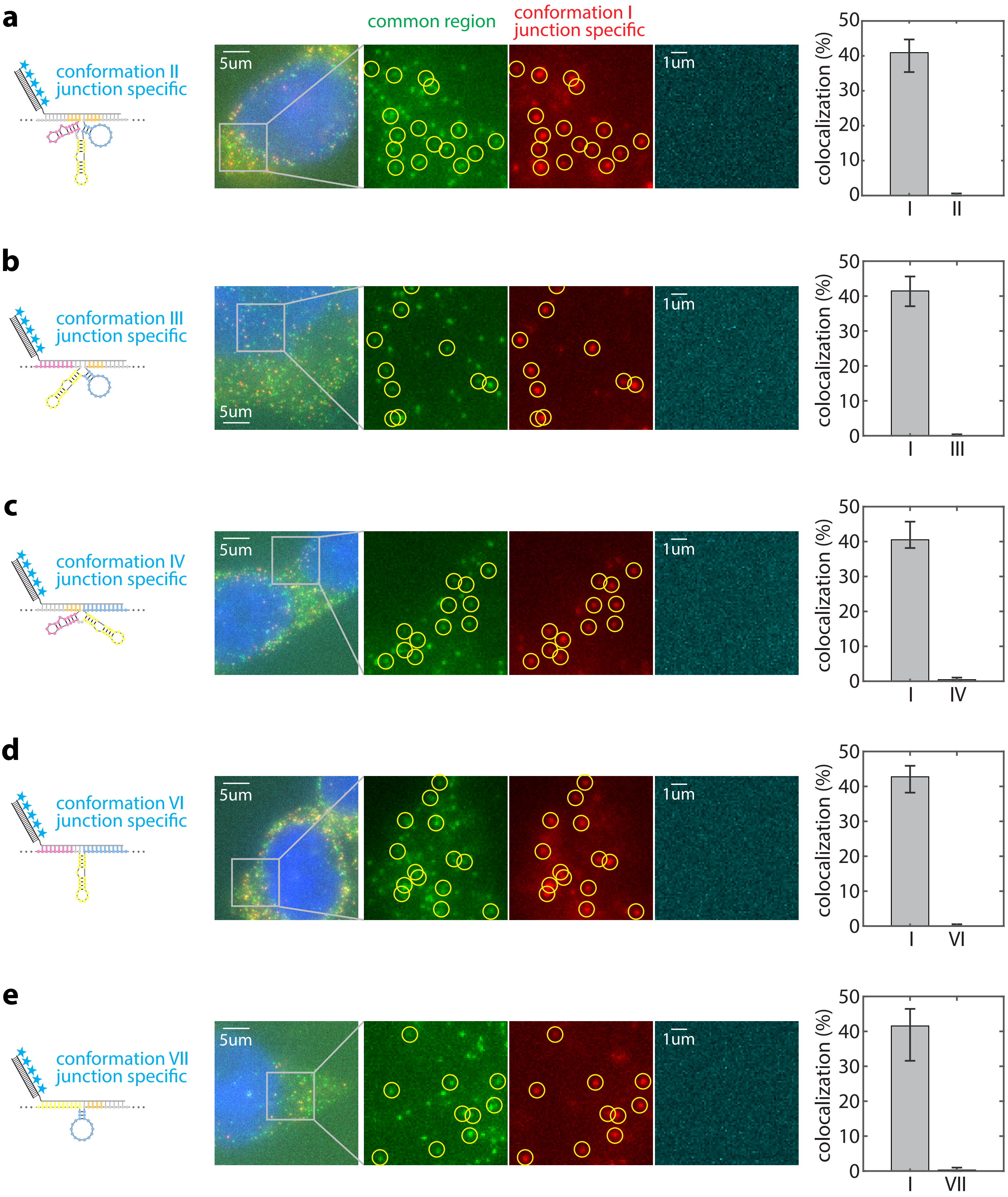

a
b
c
d
e
conformation II
junction specific
conformation III
junction specific
conformation IV
junction specific
conformation VI
junction specific
conformation VII
junction specific
common region
conformation I
junction specific
5um
1um
colocalization (%)
0
10
20
30
40
50
I
II
III
IV
VI
VII

**Supplementary Figure 13. SMOOTHY-FISH 2nd version with an initiator achieved high yield and specificity.**

**a**, In SMOOTHY-FISH 2nd version, a SMOOTHY-FISH probe is bridged to the target through the binding of an initiator (blue). We optimized a set of sequences (orange) to ensure high specificity and hybridization efficiency between the probe and the initiator. **b**, The SMOOTHY-FISH probe could not bind to the target to generate any signals ① without the initiator. **c**, SMOOTHY-FISH 2nd version showed high signal colocalization to smFISH when mapping SDHA mRNA site P26 ③ with and ② without flankers. Colocalized dots are circled in yellow; non-colocalized dots are circled in blue. **d**, SMOOTHY-FISH 2nd version approached the detection yield of the smFISH benchmark. Detection yields are shown as mean ± SD (50 cells). The benchmark is shown as mean (dash line) ± SD (gray zone).

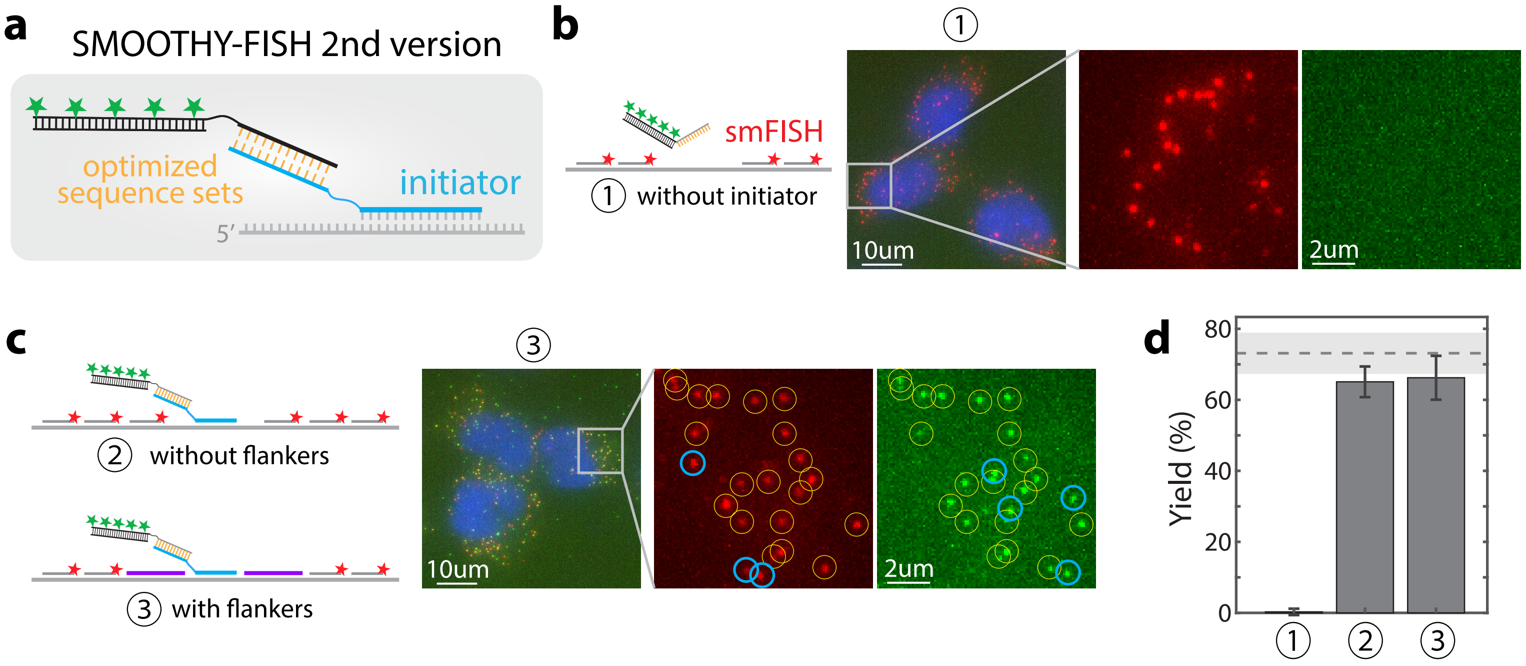
a
SMOOTHY-FISH 2nd version
optimized
sequence sets
initiator
5'
b
①
smFISH
① without initiator
10um
2um
c
③
② without flankers
③ with flankers
10um
2um
d
Yield (%)
80
60
40
20
0
①
②
③
